# ERAD Activity Distinguishes the Functional Heterogeneity of Hematopoietic Stem Cells

**DOI:** 10.64898/2025.12.01.691501

**Authors:** Qinlu Peng, Manxi Zheng, Suxuan Liu, Xinshu Xie, Zhejuan Shen, Ting Wang, Hanqi Liu, Wenbin Mi, Jie Zhou, Xuezhen Ma, Zhenyou Yin, Yuhan Hu, Guojun Shi, Yewei Ji, Dingxiao Zhang, JinHai Zheng, Xiang Wang, Kaosheng Lv, Qing Li, Jing Feng, Yang Mei, Lu Liu

## Abstract

Hematopoietic stem cells (HSCs) rely on precisely controlled proteostasis to sustain lifelong self-renewal, yet whether intrinsic proteostatic characteristics can prospectively define functional diversity within the HSC pool remains unknown. In this study, we developed a dual-fluorescence reporter system to visualize endoplasmic reticulum-associated degradation (ERAD) activity *in vivo* and discovered that ERAD activity can prospectively predicts the long-term reconstitution potential of HSCs. Remarkably, ERAD states were maintained through self-renewal, revealing an inheritable “proteostatic memory” within the HSC pool. Transcriptomic and functional analyses identified SOCS2 as a key regulator of this process: SOCS2 inhibits JAK2 signaling to sustain ERAD activity, whereas hyperactive JAK2 interacts with the ERAD E3 ligase Hrd1 and suppresses VCP-mediated misfolded substrate. These findings define a SOCS2-JAK2-ERAD axis that connects cytokine signaling to intrinsic proteostasis remodeling, establishing ERAD activity as a predictive and heritable marker of HSC function and providing a conceptual framework for understanding hematopoietic stem cell heterogeneity.

## Introduction

Hematopoietic stem cells (HSCs) sustain lifelong blood production through tightly balanced self-renewal and differentiation(*1-3*). Although phenotypically defined HSCs have long been considered a homogeneous population(*4*), recent accumulating evidence reveals marked functional heterogeneity in their lineage output, proliferative dynamics, and stress responses(*5-10*). The molecular basis of this heterogeneity remains elusive. While transcriptional, epigenetic, and metabolic cues have been implicated, few intrinsic features can prospectively distinguish HSCs with long-term regenerative potential from those destined for exhaustion(*9, 11*).

Protein homeostasis (proteostasis)-the coordinated control of protein synthesis, folding, and degradation-is essential for stem cell identity and longevity(*12*). Disruption of proteostasis drives HSC exhaustion, biased differentiation, and malignant transformation(*13-16*). Yet, whether proteostatic capacity itself constitutes a stable, heritable determinant of stem cell function remains unclear. Autophagy(*17, 18*) and the ubiquitin-proteasome system(*19*) have been shown to maintain HSC integrity, but these pathways fluctuate dynamically in response to stress and do not reliably define persistent functional states(*20*). This raises the question that whether any proteostatic characteristics could serve as an inheritable marker of HSC fitness.

The endoplasmic reticulum (ER) serves as the central compartment for the synthesis and quality control of secretory and membrane proteins. ER-associated degradation (ERAD) selectively identifies and eliminates misfolded proteins, thereby maintaining ER proteostasis under both basal and stress conditions(*21, 22*). Our previous work demonstrated that genetic disruption of the Sel1L-Hrd1 ERAD complex compromises HSC maintenance(*23, 24*), suggesting that ERAD activity represents a key determinant of stem cell fitness. However, the extent to which ERAD activity varies among individual HSCs and whether such variation reflects transient states or stable, heritable traits-have not been addressed, largely due to the absence of tools for monitoring ERAD *in vivo*.

In this study, we engineered a dual-fluorescence reporter enabling real-time visualization of ERAD activity *in vivo*. By integrating this tool with functional and transcriptomic analyses, we set out to determine whether proteostatic capacity represents an intrinsic and potentially heritable property of HSCs.

## Results

### Generation and Validation of an ERAD Activity Reporter *In Vitro* and *In Vivo*

To monitor endoplasmic reticulum-associated degradation (ERAD) (**fig. S1A**) activity in HSC *in vivo*, we developed a dual-fluorescence reporter construct, ZsGreen-P2A-NHK-DsRed Express 2. This design incorporates a self-cleaving P2A peptide (*25*) to ensure equimolar translation of ZsGreen and NHK, a well-characterized misfolded variant of α1-antitrypsin (Null Hong Kong mutant), a canonical substrate of the Sel1L-Hrd1 ERAD complex(*26, 27*) (**Fig. 1A**). Therefore, the DsRed/ZsGreen fluorescence ratio reflects ERAD-mediated degradation efficiency, independent of global protein translation, and enables real-time, chase-free quantification of ERAD activity. We validated the reporter’s reliability *in vitro* using lentiviral constructs expressing this reporter in *ER-Cre^+^; Sel1l^fl/fl^*mouse embryonic fibroblasts (MEFs)(*28*)(**fig. S1B**). Flow cytometric analysis revealed that the ZsGreen/NHK-DsRed Express 2 ratio was significantly reduced in ERAD-deficient cells compared with controls (**Fig. 1B**, **fig. S1C-D**), indicating diminished ERAD activity. Conversely, overexpression of Sel1L isoforms markedly increased the fluorescence ratio, confirming that the reporter accurately tracks ERAD alterations in murine cells (**Fig. 1C**).

**Fig. 1.**
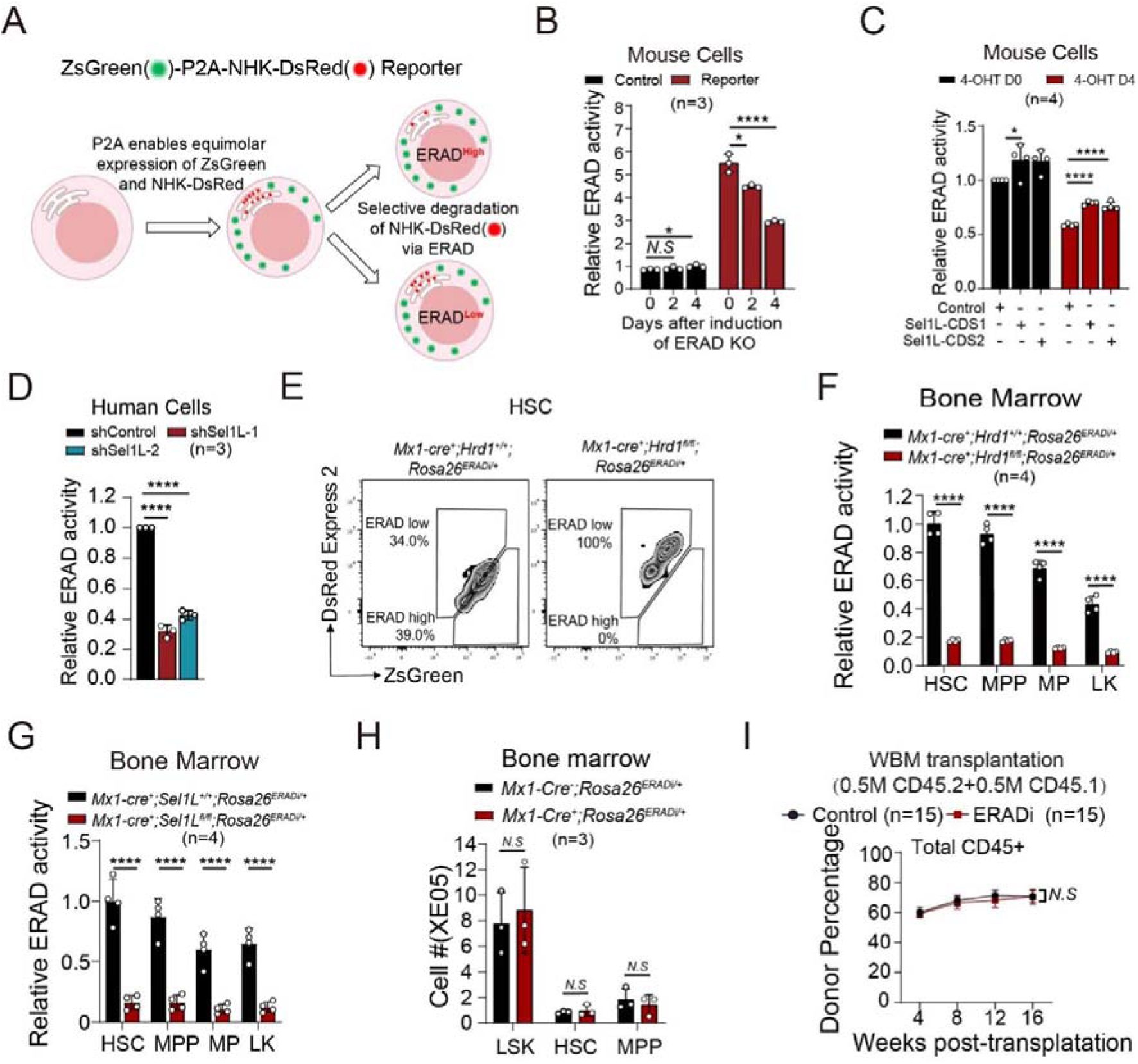
Development and validation of an *in vivo* reporter for ERAD activity. **(A)** Schematic representation of the ZsGreen-P2A-NHK-DsRed reporter construct used to assess ER-associated degradation (ERAD) activity. **(B)** *ER-Cre; Sel1L^fl/fl^* MEFs were infected with PEZ-NHK or PEZ-control lentiviral constructs and treated with tamoxifen (2Lμg/ml) for 0-4 days to induce Sel1L deletion, ERAD activity was evaluated by flow cytometry. **(C)** *ER-Cre; Sel1L^fl/fl^* MEFs were first infected with Sel1L-CDS1, Sel1L-CDS2, or control lentivirus and selected with blasticidin for 48 hours, followed by infection with PEZ-NHK or PEZ-control constructs. Cells were treated with tamoxifen (2Lμg/mL) for 0–4 days to induce Sel1L deletion, and ERAD activity was evaluated by flow cytometry. **(D)** MOLM-13 cells were infected with shSel1L or control lentivirus, selected with puromycin for 48 hours, and subjected to flow cytometric analysis of ERAD activity. **(E-F)** 6-8 weeks old *Mx1-Cre^+^; Hrd1^fl/fl^; Rosa26^ERADi/+^*and *Mx1-Cre^+^; Hrd1^+/+^; Rosa26^ERADi/+^* control mice were injected with poly(I:C) every other day for a total of three doses. Two weeks post-injection, representative flow cytometry plot showing ERAD activity in HSCs **(E)**, ERAD activity in HSPCs **(F)** were analyzed. **(G)** ERAD activity in HSPCs from *Mx1-Cre^+^; Sel1L^fl/fl^; Rosa26^ERADi/+^*and *Mx1-Cre^+^; Sel1L^+/+^; Rosa26^ERADi/+^* control mice were evaluated by flow cytometry. **(H)** *Mx1-Cre^-^; Rosa26^ERADi/+^* and *Mx1-Cre^+^; Rosa26^ERADi/+^* mice (6-8 weeks old) were injected with poly(I:C) as described above. Two weeks post-injection, the numbers of hematopoietic stem and progenitor cells were quantified. **(I)** A total of 5×10^5^ CD45.2^+^ whole bone marrow cells from *Mx1-Cre^+^; Rosa26^ERADi/+^*and control (+/+) mice transplanted together with 5×10^5^ CD45.1^+^ wild type bone marrow cells into lethally irradiated CD45.1^+^ wild-type recipients. The contribution of CD45.2 cells in total CD45^+^ cells from peripheral blood was analyzed every 4 weeks for 16 weeks. Data represent mean±s.d. from at least three independent experiments. Statistical significance was determined using unpaired two-tailed Student’s t-test.

To extend the validation, we transduced human cell line with our reporter and knocked down ERAD key components (**fig. S2A**). Knockdown of Sel1L, NPL4, and VCP all significantly impaired ERAD activity as indicated by the reporter, confirming its applicability across species (**Fig. 1D**, **fig. S1A**, **fig. S2A-B**). Furthermore, pharmacological inhibition of ERAD using NMS-873, a selective VCP inhibitor acting downstream of Sel1L-Hrd1(*29*), led to a pronounced reduction in ERAD reporter activity, under both steady-state and cycloheximide (CHX)-treated conditions (**fig. S2C**).

Based on these *in vitro* results, we generated an inducible ERAD reporter mouse strain (ERAD indicator, ROSA26^ERADi/+^) by targeting the ROSA26 locus with a floxed-stop cassette upstream of the ZsGreen-P2A-NHK-DsRed Express 2 (**fig. S3A**). Upon crossing with Mx1-Cre(*30*) transgenic mice and administering polyinosinic:polycytidylic acid (pIpC, 2.65 μg/g body weight, every other day for three doses), Cre-mediated excision was confirmed by genotyping (**fig. S3B**). Flow cytometric analysis of bone marrow cells demonstrated robust expression of both ZsGreen and DsRed Express 2 in bone marrow cells (**fig. S3C**), indicating successful hematopoietic-specific induction of the ERAD reporter *in vivo*.

### ERAD reporter Accurately Reflects Genetic Perturbations of ERAD Activity *In Vivo*

To evaluate the reliability and sensitivity of the ERAD reporter *in vivo*, we genetically disrupted core components of the ERAD machinery and assessed reporter activity across hematopoietic populations. We crossed *Mx1-Cre^+^; ROSA26^ERADi/+^* mice with *Hrd1^fl/fl^* animals*(31)* to generate *Mx1-Cre^+^; Hrd1^fl/fl^; ROSA26^ERADi/+^*mice. Following Mx1-Cre induction, flow cytometric analysis revealed a robust reduction in ERAD activity, as indicated by a decreased ZsGreen/NHK-DsRed fluorescence ratio, across all analyzed hematopoietic subsets including HSCs (Lin^-^c-Kit^+^Sca-1^+^CD48^-^CD150^+^)(*32*), multipotent progenitors (MPPs; Lin^-^c-Kit^+^Sca-1^+^CD48^-^CD150^-^), multipipe progenitors (MPs; Lin^-^c-Kit^+^Sca-1^+^CD48^+^), and Lin^-^c-Kit^+^Sca-1^-^ (LK, progenitors) populations (**Fig. 1E-F**).

As Sel1L encodes the ER-resident adaptor essential for Hrd1 function, to substantiate these findings with Hrd1 KO, we crossed *Mx1-Cre^+^; ROSA26^ERADi/+^* mice with mice harboring a floxed allele of Sel1L(*28*) to generate *Mx1-Cre^+^; Sel1L^fl/fl^; ROSA26^ERADi/+^*. Upon Mx1-Cre induction, we observed a marked and consistent reduction in ERAD reporter activity across multiple hematopoietic stem and progenitor populations (**Fig. 1G**). These results demonstrate the sensitivity and reliability of our ERAD reporter system in detecting impaired ERAD function resulting from disruption of the Sel1L-Hrd1 complex. Furthermore, analysis of hematopoietic differentiation stages revealed that ERAD activity declined with differentiation, as evidenced by lower ZsGreen/NHK-DsRed ratios in more differentiated populations compared to HSCs/MPPs (**Fig. 1F-G**). These results demonstrate that the ERAD reporter accurately reflects genetic perturbations of the ERAD pathway *in vivo*.

### Minimal Cytotoxicity of ERAD Reporter Expression on Hematopoiesis and HSC function

To assess whether our ERAD reporter induces proteotoxic stress, *Mx1-Cre^+^*mice were crossed with *ROSA26^ERADi/+^* mice to enable hematopoietic-specific, inducible expression of the reporter. ERADi expression was then induced in 6- to 8-week-old *Mx1-Cre^+^; ROSA26^ERADi/+^*mice, along with sex- and age-matched Mx1-Cre^-^; ROSA26^ERADi/+^ littermate controls. Following induction, ERADi-expressing mice exhibited comparable body weight, bone marrow cellularity, spleen and thymus cellularity compared with controls (**fig. S4A-F**). Furthermore, the absolute numbers and frequencies of HSC, MPP, MP, progenitors, and lineages were indistinguishable between ERADi-expressing and control mice (**Fig. 1H**, **fig. S4G-I**), indicating that ERADi expression does not disrupt steady-state hematopoiesis.

To further validate these findings in an independent hematopoietic-specific Cre system(*33*), we generated *Vav1-Cre^+^; ROSA26^ERADi/+^* mice and compared them to *Vav1-Cre^-^; ROSA26^ERADi/+^* littermates. Consistent with the Mx1-Cre model, *Vav1-Cre^+^; ROSA26^ERADi/+^* mice displayed normal body weight, spleen and thymus weights, bone marrow cellularity, spleen cellularity and thymus cellularity, as well as equivalent frequencies and absolute numbers of HSCs, MPP, MP, progenitors, and all mature lineage cells (**fig. S4J-R**). These findings reinforce that ERADi expression does not perturb hematopoietic homeostasis.

To assess the impact of ERADi expression on HSCs function, competitive bone marrow transplantation assays were performed. Two weeks after pIpC induction, 5 × 10^5^ whole bone marrow cells from CD45.2^+^ *Mx1-Cre^+^; ROSA26^ERADi/+^* mice or their littermate controls were transplanted alongside an equal number of CD45.1^+^ wild-type competitor cells into lethally irradiated recipients. Peripheral blood chimerism was monitored every four weeks over a 16-week period. Donor-derived multilineage reconstitution-including myeloid (Mac1^+^), B cell (B220^+^), and T cell (CD3^+^) compartments—was comparable between ERADi-expressing and control groups throughout the study (**Fig. 1I**, **fig. S5A-C**). Analysis of bone marrow at week 16 confirmed similar donor contributions to HSCs and all hematopoietic subsets (**fig. S5D-E**), indicating preserved long-term repopulation capacity. Collectively, these results demonstrate that ERADi, despite depending on a misfolded protein substrate expression, does not exert detectable cytotoxic effects on hematopoiesis or HSC function.

### ERAD Activity Indicates the Reconstitution Capacity of Bone Marrow Cells

Given the essential role of the Sel1L–Hrd1 ERAD pathway in maintaining HSC function(*23, 24*), we next investigated whether ERAD activity could predict the hematopoietic reconstitution potential of bone marrow cells. To test this hypothesis, we isolated whole bone marrow cells from pIpC-induced *Mx1-Cre^+^; ROSA26^ERADi/+^*mice based on ZsGreen/DsRed fluorescence ratio, thereby distinguishing cells with relatively high or low ERAD activity (**Fig. 2A-B**). To assess the functional differences between these sorted subsets, we transplanted 2 × 10^5^ sorted ZsGreen-positive cells with either high or low ERAD activity, together with an equal number of unsorted competitor bone marrow cells from CD45.2^+^ wild-type mice, into lethally irradiated recipient mice (**Fig. 2C**). Donor-derived peripheral blood chimerism was monitored every four weeks over a total of 16 weeks. Cells exhibiting high ERAD activity demonstrated significantly superior long-term multilineage reconstitution across total live cells, myeloid (Mac1^+^), B (B220^+^), and T (CD3^+^) cell compartments compared with those with low ERAD activity (**Fig. 2D**, **fig. S6A-C**). At the 16-week endpoint, the frequency of donor-derived HSCs and mature lineage cells in the bone marrow remained markedly higher for recipients of cells with high ERAD activity, whereas chimerism from cells with low ERAD activity progressively declined (**Fig. 2E**, **fig. S6D**).

**Fig. 2.**
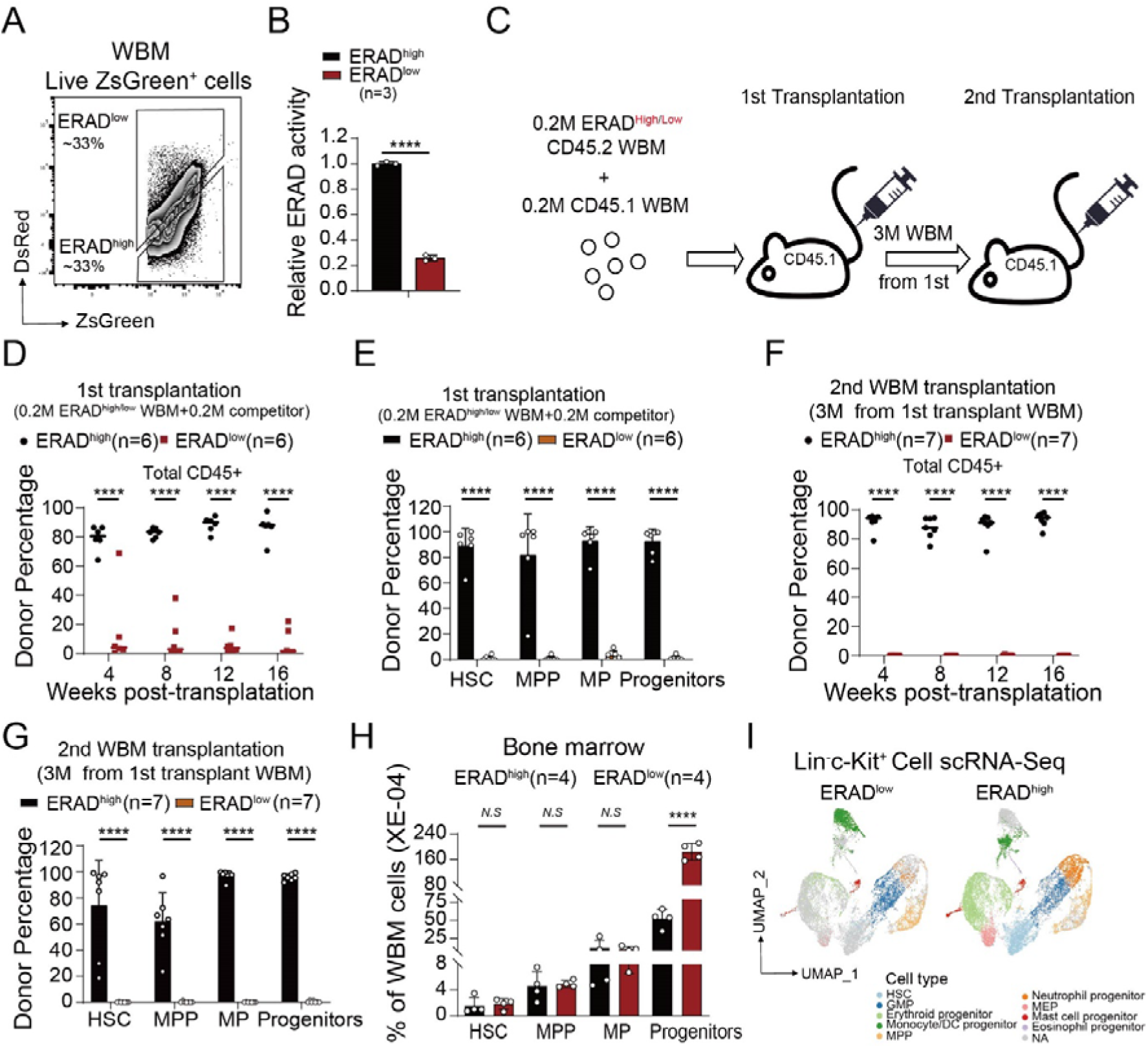
ERAD activity prospectively predicts hematopoietic stem cell reconstitution potential. **(A)** Flow cytometry gating strategy for sorting whole bone marrow cells based on ERAD activity from *Mx1-Cre^+^; Rosa26^ERADi/+^* mice following poly(I:C) induction. **(B)** Relative ERAD activity of sorted ERAD^high^ and ERAD^low^ donor bone marrow cells prior to transplantation. **(C)** Schematic of transplantation strategy. *Mx1-Cre^+^; Rosa26^ERADi/+^* mice (6-8 weeks old) were injected with poly(I:C) as described above. Two weeks post-induction, 2L×L10^5^ CD45.2^+^ ERAD^high^ or ERAD^low^ bone marrow cells were sorted and transplanted together with 2L×L10^5^ CD45.1^+^ wild-type competitor cells into lethally irradiated CD45.1^+^ recipients. For secondary transplantation, 3L×10^6^ whole bone marrow cells from primary recipients were transferred into lethally irradiated secondary CD45.1^+^ hosts. **(D-G)** Peripheral blood chimerism was assessed every 4 weeks for 16 weeks in primary **(D)** and secondary **(F)** recipients based on the contribution of CD45.2^+^ cells to total CD45^+^ cells. Contributions of CD45.2^+^ cells in HSPC compartment in the bone marrow was analyzed at the end of 16 weeks in primary **(E)** and secondary **(G)** recipients. **(H)** The proportion of hematopoietic stem and progenitor cells (HSPCs) was analyzed by flow cytometry in ERAD^high^ or ERAD^low^ whole bone marrow. **(I)** Uniform manifold approximation and projection (UMAP) plot showing cell type distributions identified by scRNA-seq in ERAD^high^ or ERAD^low^ Lineage^-^c-Kit^+^ cells. Data represent mean±s.d. from three independent experiments. Statistical significance was determined using unpaired two-tailed Student’s t-test.

To evaluate differences in long-term self-renewal capacity, we performed secondary transplantation assays by transferring bone marrow cells from primary recipient mice into secondary lethally irradiated hosts. Cells originating from the high ERAD subset not only maintained but also expanded their lineage chimerism in peripheral blood in secondary recipients (**Fig. 2F**, **fig. S6E-G**), suggesting sustained self-renewal potential. Furthermore, cells derived from the low ERAD subset showed minimal bone marrow chimerism at the experimental endpoint (**Fig. 2G**, **fig. S6H**).

Importantly, these observed functional differences in reconstitution potential were not simply attributable to differences in the initial frequency of phenotypically defined HSC, MPP, or MP within the sorted populations prior to transplantation. In fact, flow cytometric analysis revealed that the low ERAD subset unexpectedly contained a higher proportion of lineage^-^c-Kit+Sca-1^-^ cells (progenitors) (**Fig. 2H**). To address whether these flow cytometric profiles accurately reflect intrinsic functional differences, we performed single-cell RNA sequencing analysis of sorted lineage^-^c-Kit^+^ cells with high versus low ERAD activity. Given that the lineage^-^c-Kit^+^ fraction includes HSC, MPP, MP, and lineage^-^c-Kit^+^ Sca-1^-^(progenitors) cells, we reasoned that if the low ERAD subset was indeed dominated by lineage-negative c-Kit-positive Sca-1-negative progenitors, the frequencies of transcriptionally defined HSC, MPP, and MP should accordingly be lower compared to the high ERAD subset. Consistent with our expectations, single-cell RNA sequencing confirmed a substantial depletion of transcriptionally defined HSC within the low ERAD lineage^-^c-Kit^+^ subset (**Fig. 2I**), confirming that ERAD activity itself is indicative of intrinsic functional capacity rather than merely reflecting HSC abundance. These results suggest that ERAD activity can identify bone marrow subsets with superior multilineage reconstitution and self-renewal potential, independent of phenotypic definitions.

### ERAD Activity Predicts Functional Capacity of HSC Subsets

To directly assess whether ERAD activity levels within individual HSCs predict their functional potential, we performed transplantation assays using highly purified HSC subsets. One hundred fluorescence-activated cell sorting (FACS)-purified ERAD^high^ or ERAD^low^ HSCs, isolated from pIpC-induced *Mx1-Cre^+^; ROSA26^ERADi/+^*mice, were transplanted into lethally irradiated recipients together with 5 × 10^5^ CD45.2^+^ wild-type whole bone marrow competitor cells. Peripheral blood analyses revealed that ERAD^high^ HSCs contributed to much higher long-term multilineage hematopoietic reconstitution, with sustained donor chimerism across overall live cells, myeloid (Mac-1^+^), B (B220^+^), and T (CD3^+^) compartments over time (**Fig. 3A**, **fig. S7A-C**). Analysis of bone marrow at 16 weeks post-transplantation further confirmed that ERAD^high^-derived cells maintained a pool of functional HSCs and efficiently generated downstream hematopoietic progenitors and mature blood cells, while ERAD^low^-derived cells were nearly absent from these compartments (**Fig. 3B**, **fig. S7D**).

**Fig. 3.**
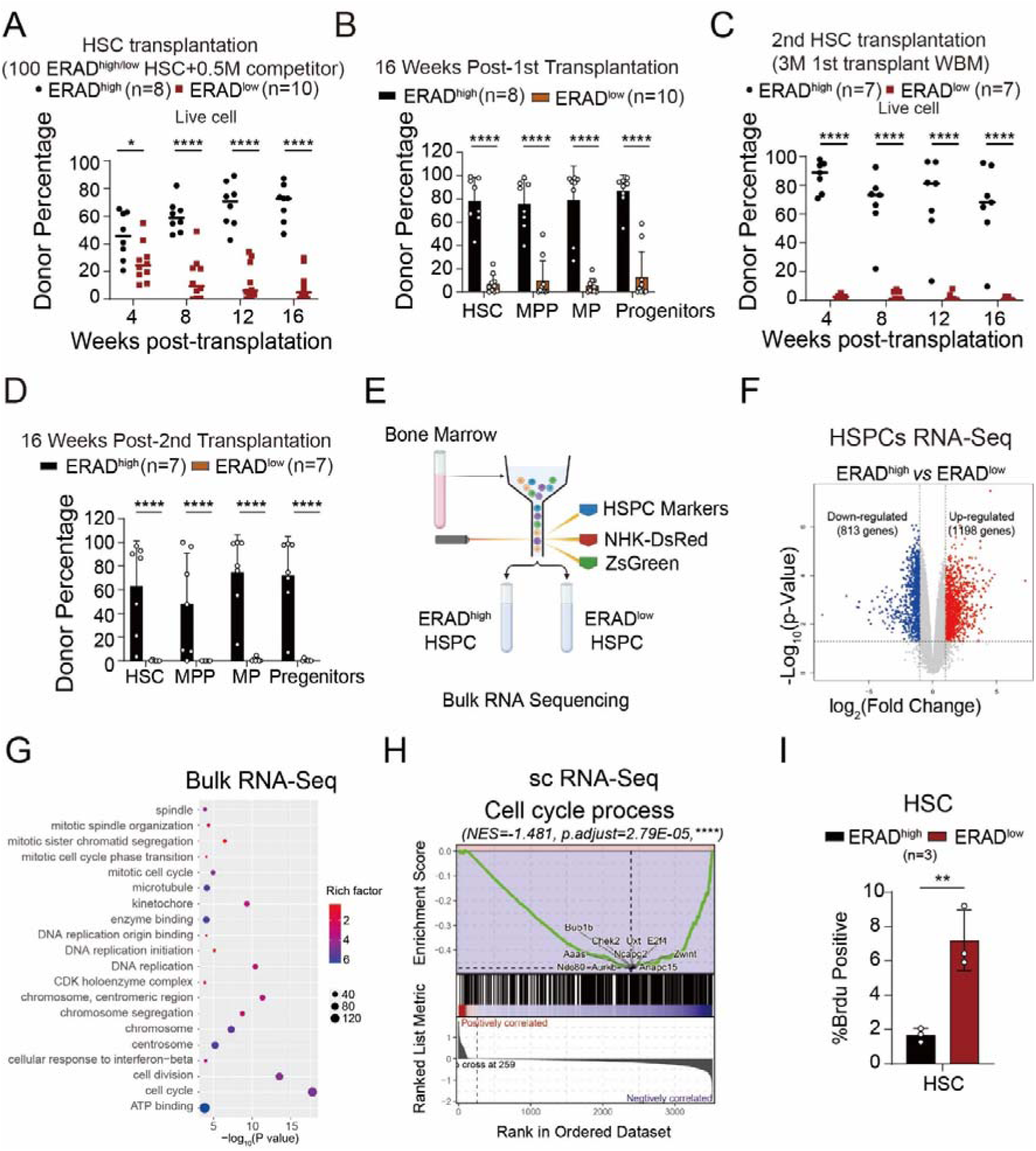
High ERAD activity defines a quiescent and functionally competent HSC state. *Mx1-Cre^+^; Rosa26^ERADi/+^* mice (6-8 weeks old) were injected with poly(I:C) as described above. **(A)** FACS-purified ERAD^high^ and ERAD^low^ ZsGreen^+^ HSCs from *Mx1-Cre^+^; Rosa26^ERADi/+^* mice were transplanted into irradiated receipt mice along with 0.5 million CD45.2 whole bone marrow competitors. **(A-D)** Peripheral blood chimerism was assessed every 4 weeks for 16 weeks in primary **(A)** and secondary **(B)** recipients based on the contribution of ZsGreen^+^ cells to total CD45^+^ cells. Contributions of ZsGreen^+^ cells in HSPC compartment in the bone marrow was analyzed at the end of 16 weeks in primary **(C)** and secondary **(D)** recipients. **(E)** Lineage^-^ c-Kit^+^ CD48^-^ HSPCs from *Mx1-Cre^+^; Rosa26^ERADi/+^*mice were sorted based on ZsGreen and DsRed fluorescence to obtain ERAD^high^ and ERAD^low^ populations for bulk RNA-seq. **(F)** Volcano map of differentially expressed genes betweenERAD^high^ and ERAD^low^ HSPCs. **(G)** GO enrichment plot of differentially expressed genes between ERAD^high^ and ERAD^low^ groups in bulk RNA-Seq data. **(H)** Gene set enrichment analysis (GSEA) plot of the cell cycle process in the differential gene enrichment analysis between ERAD^high^ and ERAD^low^ groups in single-cell RNA-sequencing data. **(I)** Two weeks after poly(I:C) injection, *Mx1-cre^+^; Rosa26^ERADi/+^* mice were injected with BrdU (200mg/kg body mass; i.p.) and then placed on drinking water containing BrdU (1 mg/ml) for 24 hours. BrdU incorporation was assessed by flow cytometry in ERAD^high^ and ERAD^low^ HSCs. Data represent mean±s.d. from three independent experiments. Statistical significance was determined using unpaired two-tailed Student’s t-test.

Secondary transplantations were performed to further evaluate long term reconstitution capacity, ERAD^high^ HSC-derived cells maintained their chimerism in secondary recipients while ERAD^low^ HSC-derived cells completely lost their reconstitution (**Fig. 3C**, **fig. S7E-G**). Again, analysis of bone marrow at 16 weeks post-transplantation further confirmed that ERAD^high^-derived cells maintained a pool of HSC, MPP, MP and progenitors, along with all mature lineage cells in bone marrow of the recipients, while ERAD^low^-derived cells were absent from all these compartments at 16 weeks post-transplantation (**Fig. 3D, fig. S7H**). Consistent with the Mx1-Cre model, Vav1-Cre–driven ERADi expression similarly confirmed that ERAD^high^ HSCs have much higher reconstitution capacity and self-renewal potential (**fig. S8A-F**). These results demonstrate that ERAD activity at the HSC level identifies functional HSCs with robust self-renewal and long-term multilineage reconstitution capacity, establishing ERAD activity as a predictive marker of HSC function.

### High ERAD Activity Marks Quiescent Hematopoietic Stem Cells

Given the strong association observed between high ERAD activity and enhanced HSC reconstitution capacity, we next sought to investigate the molecular signatures underlying this functional correlation. To address this question, we performed RNA sequencing analysis of FACS-purified hematopoietic stem and progenitor cells (HSPCs, LSK) isolated from pIpC-induced *Mx1-Cre^+^; ROSA26^ERADi/+^* mice, separated according to their relative ERAD activity levels (**Fig. 3E**). Comparative transcriptomic analysis between populations with high and low ERAD activity revealed extensive differential gene expression, identifying 1,198 significantly upregulated and 813 significantly downregulated genes associated with high ERAD activity cells (**Fig. 3F**). Notably, populations exhibiting elevated ERAD activity were enriched for transcriptional signatures previously linked to cell cycle and related programs (**Fig. 3G**). Consistently, analysis of our single-cell RNA sequencing data confirmed ERAD activity is tightly linked to cell cycle (**Fig. 3H**). Furthermore, genes closely associated with enhanced hematopoietic reconstitution capacity, including Hoxb5(*9*) (**fig. S9A-D**) and HSC dormancy genes such as Neo1(*11*), were highly enriched in ERAD^high^ HSPCs (**fig. S9A-D**).

To experimentally substantiate this correlation between ERAD activity and HSC quiescence, we performed bromodeoxyuridine (BrdU) incorporation assays to assess proliferative activity of ERAD^high/low^ HSCs. Flow cytometric analysis revealed that HSCs within the ERAD-high subset exhibited a significantly lower proportion of BrdU-positive cells compared to those with low ERAD activity (**Fig. 3I**). To investigate whether ERAD activity is accordingly altered under conditions when HSCs exit quiescence(*34, 35*), we exposed mice to acute inflammatory stress by administering high-dose poly(I:C) (10 μg/g, analyzed 48 hours post-injection) or lipopolysaccharide (2 μg/g, analyzed 24 hours post-injection). In both models, ERAD activity was markedly reduced within HSCs (**fig. S9E**). These findings suggest that elevated ERAD activity marks a quiescent and functionally competent HSC state, and that this proteostatic state is highly sensitive to inflammatory stressors that disrupt stem cell quiescence.

### Identification of Endogenous Cell Surface Markers Reflecting ERAD Activity in HSCs

Although the ERAD reporter enables precise dissection of proteostatic states, its reliance on the genetic strain limits translational application and isolation of HSCs in native settings. To enable isolation of HSCs with distinct ERAD activity independent of our genetic reporter systems, we sought to identify endogenous cell surface markers whose expression reliably correlates with intracellular proteostatic states. We hypothesized that transcriptional profiling of HSPC subpopulations with divergent ERAD activities might reveal surface proteins indicative of these internal states. To test this, we reanalyzed our bulk RNA-seq datasets and selected candidates based on stringent criteria: (1) expression fold change >2 with an adjusted p-value <0.01; (2) experimentally validated or bioinformatically predicted surface localization; (3) availability of reliable antibodies for flow cytometric validation; and (4) no prior report linking the marker to ERAD activity or HSC function (**fig. S10A**). This screening identified two novel candidates, CD200 and CD86, whose expression specifically associated with ERAD activity levels (**Fig. 4A-B**). The expression patterns of both genes were validated by RT-qPCR and scRNA-Seq (**fig. S10B-G**), and flow cytometric analysis further confirmed that CD200 (**Fig. 4C–D**) and CD86 (**Fig. 4E–F**) populations were enriched in HSPCs exhibiting high reporter-defined ERAD activity.

**Fig. 4.**
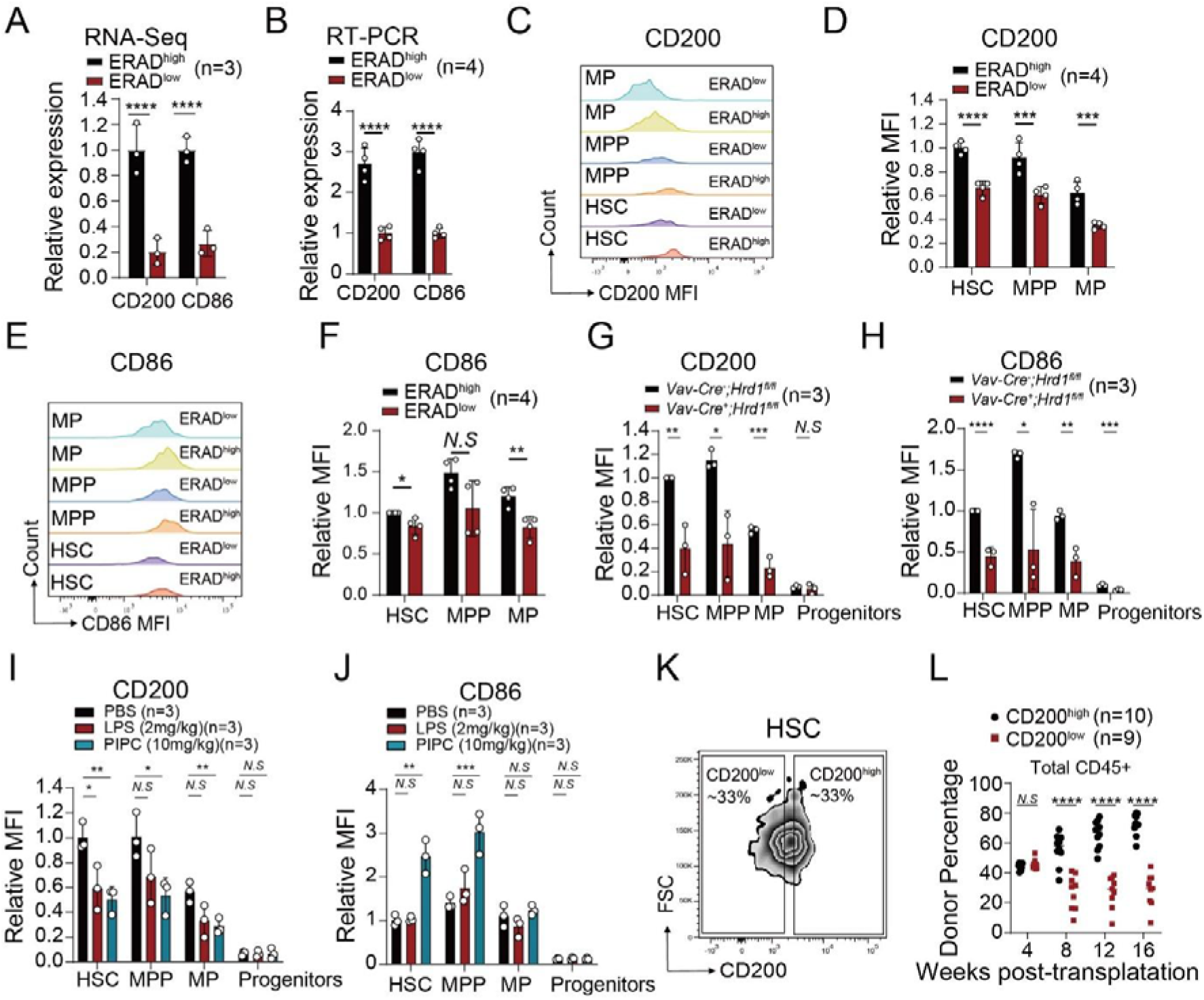
CD200 serves as an endogenous marker faithfully reflecting ERAD activity and HSC function. **(A-B)** mRNA expression of CD200 and CD86 in ERAD^high^ and ERAD^low^ HSPCs measured by bulk RNA-seq **(A)**, and validated by qRT-PCR **(B)**. **(C-F)** Representative fluorescence histogram showing CD200 **(C)** and CD86 **(E)** expression (MFI) in hematopoietic stem and progenitor cells (HSPCs) from ERAD^high^ and ERAD^low^ populations. Relative CD200 **(D)** and CD86 **(F)** expression (MFI) in HSPCs were assessed by flow cytometry from ERAD^high^ and ERAD^low^ populations. **(G-H)** CD200 **(G)** and CD86 **(H)** expression (MFI) in HSPCs from *Vav1-Cre^-^; Hrd1^fl/fl^; Rosa26^ERADi/+^* and *Vav1-Cre^+^; Hrd1^fl/fl^; Rosa26^ERADi/+^* control mice were evaluated by flow cytometry. **(I-J)** Two weeks after poly(I:C) injection, *Mx1-cre^+^; Rosa26^ERADi/+^* mice were injected with either LPS (2Lmg/kg; i.p.; 24Lhours prior to analysis) or poly(I:C) (10Lmg/kg; i.p.; 48 hours prior to analysis). CD200 **(I)** and CD86 **(J)** expression (MFI) in HSPCs from *Vav1-Cre^-^; Hrd1^fl/fl^; Rosa26^ERADi/+^*and *Vav1-Cre^+^; Hrd1^fl/fl^; Rosa26^ERADi/+^* control mice were evaluated by flow cytometry. **(K)** Flow cytometry gating strategy for sorting CD200^high^ and CD200^low^ HSCs from wild type CD45.2 mice. 100 purified CD45.2^+^ CD200^high^ or CD200^low^ HSCs were sorted and transplanted together with 5L×L10^5^ CD45.1^+^ wild-type competitor cells into lethally irradiated CD45.1^+^ recipients. **(L)** Peripheral blood chimerism was assessed every 4 weeks for 16 weeks based on the contribution of CD45.2^+^ cells to total CD45^+^ cells. Data represent mean±s.d. from three independent experiments. Statistical significance was determined using unpaired two-tailed Student’s t-test.

To further verify whether CD200 and CD86 expression faithfully reflected dynamic changes in ERAD activity, we analyzed bone marrow cells from mice with genetic deletion of key ERAD components (Sel1L or Hrd1). Both Vav-Cre– and Mx-Cre–mediated ablation led to consistent shifts in CD200 and CD86 expression (**Fig. 4G–H**, **fig. S10H–K**). We also examined expression under inflammatory conditions, which suppress ERAD activity (**fig. S9E**). Interestingly, while CD200 expression consistently mirrored ERAD dynamics across all tested conditions (**Fig. 4I**), CD86 failed to do so following lipopolysaccharide treatment (**Fig. 4J**), suggesting context-dependent limitations. Based on these findings, CD200 was chosen as the primary marker for subsequent functional studies.

We then asked whether CD200 expression reflects functional heterogeneity within phenotypic HSCs. In competitive transplantation assays, CD200^high^ HSCs exhibited superior multilineage reconstitution compared with CD200^low^ HSCs (**Fig. 4K–L**, **fig. S10L–N**). Notably, MPPs displayed CD200 expression levels comparable to HSCs. Since previous studies have suggested that subsets of MPPs share division dynamics similar to quiescent HSCs, we asked whether CD200^high^ MPPs might retain limited long-term repopulating potential. Indeed, CD200^high^ MPPs demonstrated enhanced reconstitution of lymphoid (B and T cell) but not myeloid lineages compared with CD200^low^ MPPs, although their overall reconstitution potential remained lower than that of bona fide HSCs (**fig. S10O–R**). These results establish CD200 as an endogenous cell surface marker faithfully reflecting ERAD activity and distinguish functional subsets of HSCs with different reconstitution capacity.

### SOCS2 Expression Correlates with and Predicts Heritable ERAD Activity in HSCs

We observed that ERAD activity at a given time point predicts the long-term reconstitution capacity of HSCs. This predictive association suggests that ERAD activity may represent an intrinsic, stable state rather than a transient proteostatic response. Consistently, ERAD^low^ HSCs exhibited markedly reduced reconstitution eight weeks after transplantation (**Fig. 3A**). Within this timeframe, ERAD reporter analysis in recipient bone marrow revealed that progeny derived from ERAD^high^ and ERAD^low^ HSCs retained distinct ERAD activity, although the difference was partially attenuated compared with baseline (**Fig. 5A**, **fig. S11A-B**). These results indicate that ERAD activity can be at least partially maintained through self-renewal, forming a “proteostatic memory” in HSCs.

**Fig. 5.**
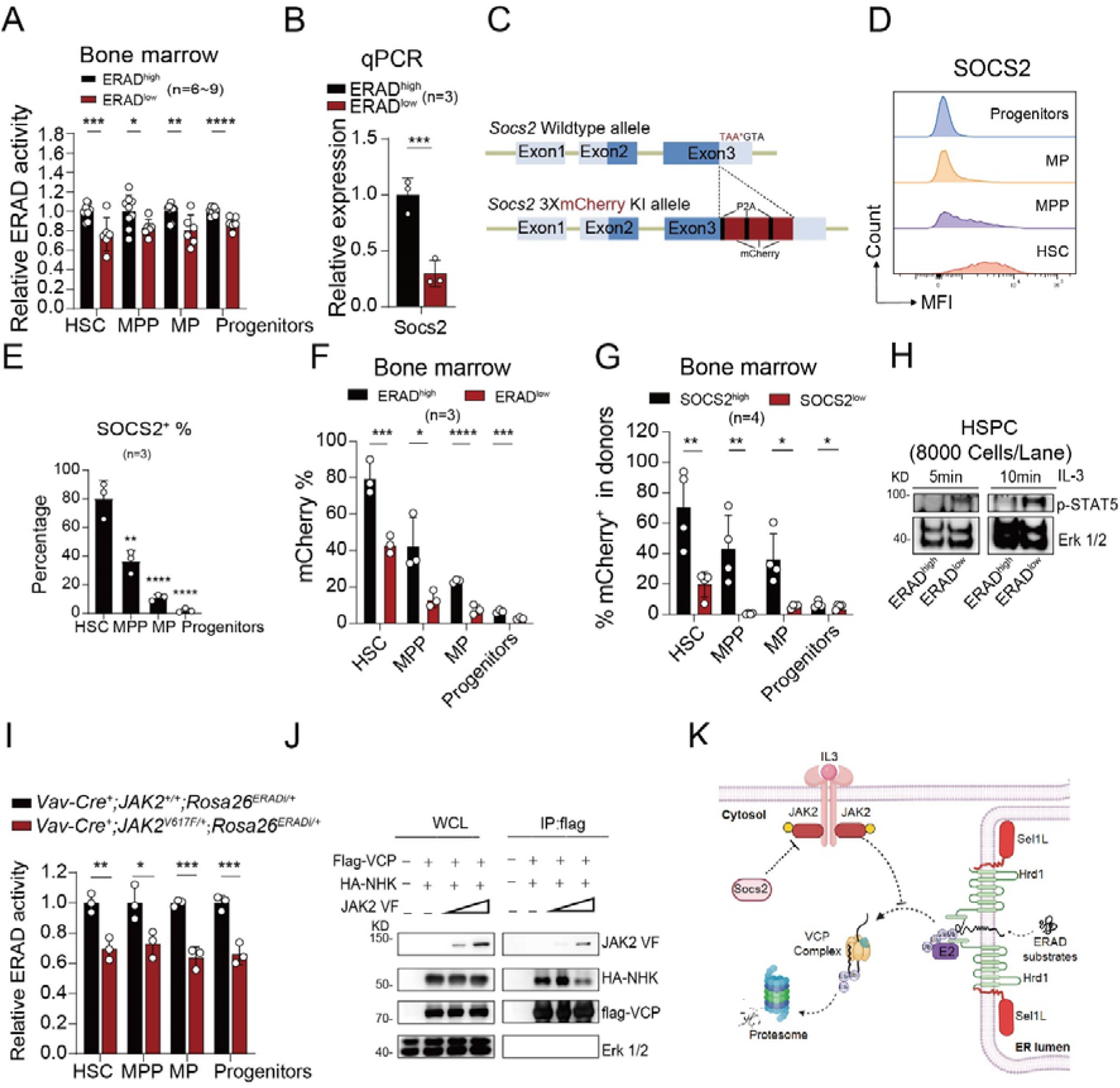
SOCS2-JAK2 signaling directly regulates ERAD activity and establishes heritable proteostatic heterogeneity in HSCs. **(A)** 100 FACS-purified ERAD^high^ and ERAD^low^ ZsGreen^+^ HSCs from *Mx1-Cre^+^; Rosa26^ERADi/+^* mice were transplanted into irradiated receipt mice along with 0.3 million CD45.2 whole bone marrow competitors. The relative ERAD activity in donor-derived hematopietic stem and progenitor cells in the bone marrow was analyzed 8 weeks post-transplantation. **(B)** mRNA expression of SOCS2 in ERAD^high^ and ERAD^low^ HSPCs was detected by qRT-PCR. **(C)** Generation strategy of the P2A-based SOCS2-3×mCherry knock-in mouse line. A 3×mCherry cassette was inserted downstream of the SOCS2 coding sequence (CDS) *via* a P2A self-cleaving peptide sequence to enable co-expression of SOCS2 and the fluorescent reporter without disrupting endogenous SOCS2 function. **(D)** Representative fluorescence histogram showing SOCS2 expression (MFI) in hematopoietic stem and progenitor cells (HSPCs). **(E-F)** Percentage of SOCS2^+^ cells in different HSPC subsets **(E)** and the mCherry proportion **(F)** in ERAD^high^ and ERAD^low^ hematopoietic stem and progenitor cells from *Mx1-Cre^+^; Rosa26^ERADi/+^* mice were assessed by flow cytometry. **(G)** 100 FACS-purified SOCS2^high^ and SOCS2^low^ CD45.2 HSCs from SOCS2-3×mCherry mice were transplanted into irradiated receipt mice along with 0.3 million CD45.2 whole bone marrow competitors. The proportion of mCherry^+^ cells in donor-derived hematopietic stem and progenitor cells in the bone marrow was analyzed 8 weeks post-transplantation. **(H)** ERAD^high^ and ERAD^low^ HSPCs were sorted and then treated with IL-3 (10ng/mL) for 5min, 10min at 37L. p-STAT5 level was detected by western blot. **(I)** ERAD activity in 6-8 weeks old *Vav1-Cre^+^; JAK2^V617F/+^; Rosa26^ERADi/+^* and *Vav1-Cre^+^; JAK2^+/+^; Rosa26^ERADi/+^* control mice were analyzed by flow cytometry. **(J)** HEK 293T cells were transfected with constructs that expressed FLAG-tagged VCP, Myc-tagged JAK2-V617F, HA-tagged NHK. VCP proteins were immunoprecipitated using FLAG antibody, competitive binding of JAK2-V617F and NHK to VCP were detected by western blot. **(K)** Schematic model of SOCS2-JAK2 kinase signaling directly regulates ERAD activity. Data represent mean±s.d. from three independent experiments. Statistical significance was determined using unpaired two-tailed Student’s t-test.

To identify factors underlying this heritable ERAD activity diversity, we first examined whether canonical UPR signaling contributes to its regulation. Bulk RNA-seq analysis revealed no significant correlation between ERAD activity and the expression of classical UPR target genes, including those regulated by ATF6 or spliced XBP1 (**fig. S12A**). RT–qPCR analysis further confirmed comparable expression of ERAD- and UPR-related genes between ERAD^high^ and ERAD^low^ HSPCs (**fig. S12B**), indicating that basal UPR signaling may not account for the observed heterogeneity.

We next performed an integrative screen combining bulk and single-cell transcriptomics to identify candidate regulators of ERAD activity. Genes were prioritized based on (i) differential expression between ERAD^high^ and ERAD^low^ HSPCs, (ii) enrichment among the top 300 differentially expressed genes within cluster 0 in scRNA-seq (**fig. S13A-B**), and (iii) progressively downregulated during HSC differentiation(*36*). This analysis identified 12 candidates (**fig. S13C**), among which SOCS2 was selected for further study because it is a known negative regulator of JAK2 signaling and both JAK2 and ERAD influence the expression and membrane localization of the thrombopoietin receptor MPL(*37, 38*).

After confirming that Socs2 expression was significantly enriched in ERAD^high^ HSPCs by qPCR (**Fig. 5B**), we generated a knock-in SOCS2 reporter to further validate this pattern, showing a substantially higher percentage of SOCS2^+^ cells within the HSC compartment relative to committed progenitors (**Fig. 5C–E**). Notably, the SOCS2^+^ fraction was markedly higher in ERAD^high^ than in ERAD^low^ HSCs (**Fig. 5F**), establishing a strong positive correlation between SOCS2 expression and ERAD activity.

To assess whether SOCS2 expression is maintained in self-renewing progeny, we transplanted purified SOCS2^high^ and SOCS2^low^ HSC subsets from SOCS2 reporter mice. Eight weeks post-transplantation, we observed a significant difference of SOCS2 expression between progeny derived from SOCS2^high^ versus SOCS2^low^ donors (**Fig. 5G**), suggesting a heritable SOCS2 expression during HSC self-renewal, analogous to ERAD activity. (**Fig. 5A**, **fig. S11A-B**).

### SOCS2–JAK2 Signaling Directly Controls ERAD Activity in HSCs

To determine whether SOCS2 directly regulates ERAD activity, we silenced Socs2 using shRNAs, which efficiently reduced Socs2 mRNA levels and led to a marked suppression of ERAD reporter activity (**fig. S13D–E**). Given that SOCS2 functions as a classical negative feedback regulator of JAK2/STAT signaling, we next examined whether ERAD^high^ and ERAD^low^ HSPCs exhibit differential cytokine responsiveness. Immunoblotting analysis revealed that ERAD^high^ HSPCs display substantially lower IL-3–induced STAT5 phosphorylation compared with ERAD^low^ cells, whereas responses to TPO or SCF were largely comparable (**Fig. 5H**, **fig. S13F**). This pattern mirrors the hyperactivation of IL-3-JAK2-STAT5 signaling previously reported in Socs2-null hematopoietic cells(*39*), supporting the notion that SOCS2 modulates ERAD activity through JAK2 signaling. Furthermore, pharmacological inhibition of JAK2 with Fedratinib or Ruxolitinib significantly enhanced ERAD reporter activity in human leukemic cell lines (**fig. S13G–I**). Moreover, cycloheximide chase assays confirmed that Fedratinib suppressed JAK2 phosphorylation and concurrently accelerated degradation of the ERAD substrate FLAG-NHK (**fig. S13J**), suggesting a direct inhibitory effect of JAK2 signaling on ERAD function. To further substantiate these findings *in vivo*, we assessed ERAD activity in *Vav1-Cre^+^; ROSA26^ERADi/+^; JAK2^V617F/+^*mice expressing a constitutively active JAK2 mutant. Flow cytometric analysis revealed significantly reduced ERAD reporter activity in JAK2^V617F/+^ HSCs relative to littermate controls (**Fig. 5I**).

To explore the molecular basis of JAK2-mediated ERAD regulation, we performed co-immunoprecipitation assays, which showed that wild-type JAK2 physically associates with the ERAD E3 ubiquitin ligase Hrd1, and that the *JAK2 V617F* mutation markedly strengthens this interaction (**fig. S13K**). Given that both Hrd1 and JAK2 have been reported to associate with the ERAD substrate MPL and modulate its surface expression(*37, 38*), and that JAK2 can phosphorylate the AAA-ATPase VCP(*40*)-a key component responsible for substrate extraction from the ER membrane-we hypothesized that hyperactive JAK2 signaling may disrupt VCP-mediated substrate dislocation. Indeed, enforced expression of JAK2 markedly reduced the association between VCP and ERAD substrates (**Fig. 5J**).

Finally, to assess whether Socs2 expression correlates with HSC functional potential, one hundred FACS-purified Socs2^high^ or Socs2^low^ HSCs from Socs2 reporter mice were competitively transplanted with wild-type bone marrow cells. Socs2^high^ HSCs showed markedly superior long-term multilineage reconstitution, maintaining donor chimerism across myeloid, B, and T lineages (**fig. S13L–O**), consistent with the strong association between Socs2 expression, ERAD activity, and functional competence.

Together, these findings indicate that SOCS2-JAK2 kinase signaling directly restrains ERAD activity (**Fig. 5K**), thereby linking extrinsic cytokine signaling to the intrinsic proteostatic network that shapes HSC functional heterogeneity.

## Discussion

Proteostasis has been previously considered as a transient homeostatic process, yet whether it has stable and heritable features in stem cell identity was largely unexplored. Here, we demonstrate that proteostatic characteristics can function as inheritable determinants of stem cell identity, using ERAD as a model system. ERAD-mediated proteostasis emerges as a stable and heritable feature that defines functional heterogeneity within the hematopoietic stem cell (HSC) compartment. Visualization of ERAD activity *in vivo* reveals that proteostatic capacity prospectively predicts HSC potential and remains partially preserved through self-renewal, indicating a “proteostatic memory” that underlies long-term functional diversity. Whereas other proteostasis pathways such as autophagy(*20*) are transient and dynamic, ERAD activity specifically marks quiescent, long-term reconstituting HSCs, suggesting that sustained ER integrity provides a proteostatic foundation for stem cell longevity.

Mechanistically, our findings establish a SOCS2-JAK2-ERAD axis that integrates cytokine signaling with intrinsic protein quality control. SOCS2, enriched in ERAD^high^ HSPCs and maintained in their progeny, restrains JAK2 activity to sustain ERAD function, whereas hyperactive JAK2 interacts with the ERAD E3 ligase Hrd1 and disrupts VCP-mediated substrate extraction. This inverse coupling suggests that local variations in cytokine gradients or inflammatory cues can reprogram proteostatic states, generating durable functional diversity from an initially homogeneous HSC pool. Such signaling-dependent modulation of ER proteostasis may represent a physiological mechanism through which environmental inputs imprint intrinsic differences among otherwise equivalent stem cells.

The heritable nature of ERAD states implies that proteostasis programs established in HSCs can propagate along self-renewal, influencing lineage bias and proliferative behavior. This finding extends the concept of stem cell “memory” beyond transcriptional or epigenetic inheritance, highlighting proteostasis as a previously unrecognized layer of intrinsic regulation that organizes HSC heterogeneity. The dual-fluorescence ERAD reporter and CD200 surface correlate developed here further provide genetic-independent means to resolve proteostatic heterogeneity *in vivo*, offering a functional rather than purely phenotypic framework to define HSC states. Together, these results propose a proteostasis-centered model in which extrinsic cues and intrinsic protein quality control jointly shapes the landscape of stem cell diversity, advancing our understanding of how stable functional heterogeneity arises within the HSC compartment.

## Acknowledgments

We thank Analytical Instrumentation Center of Hunan University for providing for technical assistance.

## Author contributions

Conceptualization: JF, YM, LL

Methodology: TW, HL, GS, YJ, GS, D.Z, J.Z, K.L, Q.L,

Investigation: QP, MZ, SL, XX, ZS, TW, HL, WM, JZ, XM, ZY, YH

Funding acquisition: LL

Project administration: JF, YM, LL

Supervision: JF, YM, LL

Writing – original draft: QP, MZ, SL and XX

Writing – review & editing: JF, YM, LL

## Data availability

All other data supporting the findings of this study are available from the corresponding author upon reasonable request.

## Supplementary Materials

Materials and Methods

Figs. S1 to S13

Tables S1 to S3

References (*41*)

