## Supplementary Materials for "ERAD Activity Distinguishes the Functional Heterogeneity of Hematopoietic Stem Cells"

### Methods

#### Mice

All the mice used in this study were listed in **table. S1** and maintained in C57BL background. Recipients in reconstitution assays were adult C57BL/Ka-CD45.1: Thy-1.2 or C57BL/Ka-CD45.2: Thy-1.1 mice, at least 8 weeks of age at the time of irradiation. pIpC was reconstituted in PBS and administered at 2.65 μg/gram body mass/day by intraperitoneal injection. All mice were analyzed at 8-14 weeks of age, a minimum of two weeks after pIpC treatment, and paired with sex- and age-matched controls. No phenotypic differences were observed between male and female mice. An equal number of male and female mice were used for analyses when possible and were pooled together for analysis.

#### Genotyping

150 μL 50 mM NaOH was added into a 1.5 ml tube containing 1-2 mm portion of tail or at least 1 million cells of each mouse. Each sample was heated to 90 °C for 1 hours. 50 μL of Tris-Hcl (pH=8) was then added to each tube to neutralize the reaction. All the samples were then vortexed and 2 μl of each sample was for each genotyping PCR reaction. GoTaq® Green Master Mix was used for Genotyping PCR following the instructions. All the primers were listed in **table. S2**.

#### Competitive repopulation assay

Adult recipient mice (CD45.1 or CD45.2) were irradiated with the Rad Source RS2000pro, delivering approximately 0.75 Gy/min to a total dose of 7.5 Gy. Cells were injected into the tail veins of the recipients. Beginning 4 weeks after transplantation and continuing for at least 16 weeks, blood was obtained from the tail veins of the recipient mice, subjected to ammonium-chloride potassium red cell lysis buffer (8 g/L NH4Ac, 1 g/L KHCO3, 0.04 g/L EDTA), and stained with directly conjugated antibodies to CD45.2 (104), CD45.1 (A20), B220 (6B2), Mac-1 (M1/70), CD3 (KT31.1), and Gr-1 (8C5) to monitor engraftment (Lineage Bleeding Cocktail in **table. S3**). The chimerism of stem cells, progenitors, and lineage cells was analyzed at least 16 weeks after transplantation unless otherwise indicated. Additional information on antibodies is provided in the **table. S3**.

**Flow cytometry and isolation of hematopoietic cells.**

Bone marrow cells were flushed from the femurs and tibiae of mice using FACS buffer (Hank's buffered salt solution without calcium or magnesium, supplemented with 2% heat-inactivated calf serum). The collected cells were filtered through a 70 µm cell strainer to obtain single-cell suspensions. For analytical flow cytometry, cells were stained with the antibody cocktail listed in **table. S3** and analyzed using a BD FACS Fortessa cytometer. For cell sorting experiments, total bone marrow cells were harvested from all major bones and stained with the sorting antibody cocktail described in **table. S3**. After one wash, cells were incubated with anti-APC magnetic beads (Miltenyi Biotec) for 30 minutes, followed by magnetic enrichment of c-Kit$^{+}$ cells using MS columns (Miltenyi Biotec). Dead cells were excluded using DAPI (4′,6-diamidino-2-phenylindole) staining. Final cell sorting was performed on a BD FACS Aria III flow cytometer. All reagents and materials used in these experiments are listed in **table. S3**.

**Brdu Assay**

Mice were intraperitoneally injected with a single dose of BrdU (200 mg/kg body mass) and subsequently maintained on BrdU-containing (1 mg/ml) drinking water for 24 hours. BrdU incorporation was assessed using the APC BrdU Flow Kit (BD Biosciences, 557892#) according to the manufacturer's instructions. Antibody staining was performed with the cocktail listed in **table. S3**.

**Co-immunoprecipitation**

HEK293T cells were transfected with plasmids in advance. Briefly, cells were harvested and lysed using cell lysis buffer (150 mM NaCl, 1 mM EDTA, 50 mM Tris-HCl pH 7.5 or 8.0, protease and phosphatase inhibitors, 10 mM N-ethymaleimide, and 1% Nonidet P-40 (NP-40)). The lysates were incubated with anti-FLAG magnetic beads (Thermo Fisher A36798) to capture FLAG-tagged protein complexes. Immunoprecipitates were isolated using a magnetic rack, washed thoroughly, and eluted by boiling in 2×SDS loading buffer at 95 °C for 10 minutes.

**Western blotting**

For cell line samples, 200,000 cells were collected. For mouse LSK samples, 5,000 $ERAD^{high}$ or $ERAD^{low}$ LSK cells were sorted into IMDM supplemented with 10 ng/mL TPO and 1% BSA, followed by incubation at 37 °C for 2 minutes. Cells were then washed once with PBS and precipitated using trichloroacetic acid (TCA) to a final concentration of 10%. Samples were incubated at 4 °C overnight, centrifuged at 13,000 rpm for 10 minutes at 4 °C, and the resulting pellets were washed twice with pre-chilled acetone. After air-drying, the pellets were dissolved in protein solubilization buffer (9 M Urea, 2% Triton X-100, 1% DTT), mixed with SDS loading buffer, and boiled at 95 °C for 10 minutes. Proteins were separated on 4–12% precast SDS-PAGE gels and transferred to PVDF membranes. Antibodies used for immunoblotting are listed in **table. S1**.

**Lentiviral Production and Transduction of Suspension Cells assay**

Lentiviral particles were produced by co-transfecting HEK293T cells with the lentiviral expression vector along with the packaging plasmid psPAX2 and envelope plasmid pMD2.G. Viral supernatants were collected 48 hours post-transfection and filtered through a 0.45 μm filter to remove cell debris. Exponentially growing MOLM-13 cells were infected with the filtered viral supernatant supplemented with 10 μg/mL polybrene. The infection was enhanced by centrifugation at 2,500 rpm for 90 minutes at 37 °C. Cells were then incubated at 37 °C for an additional 8 hours, after which the viral supernatant was replaced with fresh culture medium. Cells were cultured for another 48 hours. Successfully transduced cells were selected either by antibiotic resistance (puromycin or blasticidin) or by sorting for fluorescent marker-positive cells via flow cytometry.

**Cycloheximide (CHX) assay**

MOLM-13 cells overexpressing PBE2-NHK-FLAG were cultured in RPMI-1640 medium supplemented with 10% fetal bovine serum (FBS) and 1% penicillin-streptomycin at 37 °C in a 5% $CO_2$ incubator, and maintained in the logarithmic growth phase. Cycloheximide (CHX) was dissolved in DMSO to prepare a 10 mg/mL stock solution. For treatment, the stock was diluted in complete medium to a final concentration of 100 μg/mL and added to the cells. A DMSO vehicle control was included in parallel. Cells were incubated with CHX for 0, 3, or 6 hours, then harvested, washed twice with PBS, and counted. Samples were subsequently used for Western blot analysis to assess NHK protein stability.

**RT-qPCR**

LSK cells were directly sorted into 500 µL of TRIzol reagent. RNA extraction was performed according to the manufacturer's instructions. During the extraction process, 12 µL of Linear Acrylamide was added to each sample to enhance RNA precipitation. The final RNA was dissolved in DEPC-treated water and reverse-transcribed into cDNA using the High-Capacity RNA-to-cDNA™ Kit (Thermo Fisher Scientific). Quantitative PCR was performed using Power SYBR™ Green PCR Master Mix (Thermo Fisher Scientific). Reagents used in the experiment are listed in **table. S1**, and primer sequences are provided in **table. S2**.

**RNA extraction, library preparation and sequencing**

Total RNAs were extracted using TRIzol Reagent (Invitrogen, cat. NO 15596026) following the methods by Chomczynski et al(*41*). DNA digestion was carried out after RNA extraction by DNaseI. RNA quality was determined by examining A260/A280 with a NanoDrop spectrophotometer (Thermo Fisher Scientific Inc). RNA Integrity was confirmed by 1.5% agarose gel electrophoresis. Qualified RNAs were finally quantified by Qubit3.0 with Qubit RNA Broad Range Assay kit (Life Technologies, Q10210). A total of 2 µg RNAs were used for stranded RNA sequencing library preparation using KC-Digital Stranded mRNA Library Prep Kit for Illumina® (Catalog NO. DR08502, Wuhan Seqhealth Co., Ltd. China) following the manufacturer's instruction. The kit eliminates duplication bias in PCR and sequencing steps, by using unique molecular identifier (UMI) of 8 random bases to label the pre-amplified cDNA molecules. The library products corresponding to 200-500 bps were enriched, quantified and finally sequenced on DNBSEQ-T7 sequencer (MGI Tech Co., Ltd. China) with PE150 model.

**RNA-Seq data analysis**

UID RNA-seq experiment and high through-put sequencing and data analysis were conducted by Seqhealth Technology Co., LTD (Wuhan, China). Raw sequencing data was first filtered by Trimmomatic (version 0.36), low-quality reads were discarded and the reads contaminated with adaptor sequences were trimmed. Clean Reads were further treated with in-house scripts to eliminate duplication bias introduced in library preparation and sequencing. In brief, clean reads were first clustered according to the UMI sequences, in which reads with the same UMI sequence were grouped into the same cluster. Reads in the same cluster were compared to each other by pairwise alignment, and then reads with sequence identity over 95% were extracted to a new sub-cluster. After all sub-clusters were generated, multiple sequence alignment was performed to get one consensus sequence for each sub-cluster. After these steps, any errors and biases introduced by PCR amplification or sequencing were eliminated.

The de-duplicated consensus sequences were used for standard RNA-seq analysis. They were mapped to the reference genome using STAR software (version 2.5.3a) with default parameters. Reads mapped to the exon regions of each gene were counted by featureCounts (Subread-1.5.1; Bioconductor), and then RPKM was calculated. Genes differentially expressed between groups were identified using the edgeR package (version 3.12.1). A p-value cutoff of 0.05 and Fold-change cutoff of 2 were used to judge the statistical significance of gene expression differences. Gene ontology (GO) analysis and Kyoto encyclopedia of genes and genomes (KEGG) enrichment analysis for differentially expressed genes were both implemented by KOBAS software (version: 2.1.1) with a P-value cutoff of 0.05 to judge statistically significant enrichment. Alternative splicing events were detected by using rMATS (version 3.2.5) with an FDR value cutoff of 0.05 and an absolute value of $\Delta\psi$ of 0.05.

**Quantification and statistical analysis**

All quantitative data are presented as mean ± standard deviation, unless stated otherwise. For statistical analysis, two-tailed Student's t-tests were employed for comparisons between two groups, while ANOVA was used for comparisons involving more than two groups, using Prism 7 (GraphPad Software), Significance thresholds were defined as*P < 0.05, **P < 0.01, ***P < 0.005, ****P < 0.001. No randomization or blinding was applied in any experiments, and no experimental mice were excluded from the analysis. Sample sizes were determined based on experimental variation within control groups and are specified in the figure legends.

Table S1. Reagents and Resources

| Reagents or Resources | Source | Identifier |
|---|---|---|
| FACS Antibodies | | |
| Anti-mouse Mac-1 | Biolegend | Clone: M1/70, Cat#: 101212 |
| Anti-mouse Gr-1 | Biolegend | Clone: RB6-8C5, Cat#:108406,108408,108404 |
| Anti-mouse B220 | Biolegend | Clone: RA3-6B2, Cat#: 103236,103206,103204, 103208 |
| Anti-mouse CD2 | Biolegend | Clone: RM2-5, Cat#: 100105,100107,100104 |
| Anti-mouse CD3 | Biolegend | Clone:17A2, Cat#:100204,100206,110304, 100312 |
| Anti-mouse CD5 | Biolegend | Clone: 53-7.3, Cat#:100606,100608,100604 |
| Anti-mouse CD8 | Biolegend | Clone: 53-6.7, Cat#:100708,100706,100704 |
| Anti-mouse Ter-119 | Biolegend | Clone: Ter-119, Cat#:116206,116208,116204 |
| Anti-mouse CD48 | Biolegend | Clone: HM48-1, Cat#:103424,103426 |
| Anti-mouse CD150 | Biolegend | Clone: TC15-12F12.2, Cat#:105904,105914 |
| Anti-mouse CD45.2 | Biolegend | Clone: A20, Cat#:110738 |
| Anti-mouse CD45.1 | Biolegend | Clone: A20, Cat#:109822 |
| Anti-mouse CD135 | Biolegend | Clone: A2F10, Cat#: 135306 |
| Anti-mouse CD16/32 | Thermo Fisher | Clone: 93, Cat#:25-0161-82 |
| Anti-mouse CD34 | Thermo Fisher | Clone: RAM34, Cat#:11-0341-82 |
| Anti-mouse c-kit | Biolegend | Clone: A7R34, Cat#: 105826, 105825 |
| Anti-mouse Sca-1 | Biolegend | Clone: D7, Cat#: 108124, 108126, 108143 |
| Anti-mouse CD200 | Biolegend | Clone: OX-104, Cat#: 123817,329207 |
| Anti-mouse CD86 | Biolegend | Clone: GL-1, Cat#: 105013 |
| Brilliant Violet 510™ Streptavidin | Biolegend | Cat#: 405234 |
| Western Blot Antibodies | | |
| Anti-rabbit IgG, HRP-linked | Cell Signaling Technology | Clone:N/A Cat#:7074 |
| Anti-mouse IgG,HRP-linked | Cell Signaling Technology | Clone:N/A Cat#:7076 |
| Phospho-Jak2 (Tyr1007/1008) Antibody | Cell Signaling Technology | Clone:N/A Cat#:3771 |
| Phospho-Stat5 (Tyr694) Antibody | Cell Signaling Technology | Clone:N/A Cat#:9351 |
| β-actin polyclonal antibody | Proteintech | Clone:N/A Cat#:20536-1-AP |
| ERK1/2 | Cell Signaling Technology | Clone:137F5 Cat# 4695 |
| Monoclonal anti-FLAG® M2 mouse antibody | Sigma-Aldrich | Clone: N/A, Cat#:F1804 |
| HA-Tag (C29F4) Rabbit mAb | Cell Signaling Technology | Clone: C29F4, Cat#:3724S |
| Myc-Tag (71D10) Rabbit mAb | Cell Signaling Technology | Clone: 71D10, Cat#:2278S |
| Chemicals, Peptides, and Recombinant Proteins | | |
| ER-Tracker™ Red | Thermo Fisher | Cat#: E34250 |
| HBSS, no calcium, no magnesium, no phenol red | Thermo Fisher | Cat#: 14175095 |
| Fetal Bovine Serum (FBS) | Sera Pure | Cat#:SE141 |
| DAPI (4',6-Diamidino-2-Phenylindole, Dilactate) | Biolegend | Cat#: 422801 |

| | | |
|---|---|---|
| Power SYBR™ Green PCR Master Mix | Thermo Fisher | Cat#: 4367659 |
| Gibco™IMDM (Iscove's Modified Dulbecco's Medium) | Thermo Fisher | Cat#: 12440053 |
| Dimethyl sulfoxide(DMSO) | Sigma Aldrich | Cat#: D2650-100ML |
| TRIzol™ Reagent | Thermo Fisher | Cat#: 15596026 |
| Chloroform | Sigma Aldrich | Cat#: C2432-500ML |
| 2-Propanol | Sigma Aldrich | Cat#:190764-500ML |
| Linear Acrylamide | Thermo Fisher | Cat#: AM9520 |
| Trichloroacetic acid | MP biomedicals | Cat#: SKU 02196057-CF |
| Urea | sangon | Cat#: A610148-0500 |
| ammonium acetate (NH4Ac) | sangon | Cat#: A501574-0500 |
| potassium bicarbonate (KHCO3) | sangon | Cat#: A501195-0500 |
| Ethylenediaminetetraacetic acid tetrasodium salt dihydrate (EDTA) | sangon | Cat#: A610185-0500 |
| Triton™ X-100 | sangon | Cat#: A600198-0500 |
| 1M Tris-HCl Solution, pH 8.0, Sterile | sangon | Cat#: B548127-0500 |
| Nonidet（R）P-40 | sangon | Cat#: A600385-0500 |
| Tris | sangon | Cat#: A600194-0500 |
| N-Ethylmaleimide | sangon | Cat#: A600450-0005 |
| Recombinant Murine SCF | PeproTech | Cat#: 250-03 |
| Recombinant Murine TPO | PeproTech | Cat#: 315-14 |
| S-Clone SF-O3 Culture Medium | Iwai North America Inc. | SKU:1303 |
| MethoCult™ GF M3434 | Stem cell technolgies | Cat#: M3434 |
| pIpC | Amersham biosciences(Now GE lifesciences) | Cat#: 27473201 |
| Omni-Easy™ Fast Protein Loading buffer (5X) | Epizyme | Cat#: LT101 |
| Cycloheximide | Sigma Aldrich | Cat#: C7698-1G |
| DEPC Water | Invitrogen Life Technologies | Cat#: 46-2224 |
| GoTaq® Green Master Mix | Promega | Cat#: M7123 |
| Sodium chloride | sangon | Cat#: A100241-0500 |
| DTT | sangon | Cat#: A620058-0005 |
| Sodium hydroxide | sangon | Cat#: A100173 |
| Blasticidin | Invitrogen | Cat#:ILTR21001 |
| Polybrene | Merck millipore | Cat#:TR-1003-G |
| Ruxolitinib | Selleck | Cat#:S1378 |
| Liberase TL | Roche | Cat#:05401020001 |
| Fedratinib | MedChemExpress | Cat#:HY-10409 |
| Fedratinib | TargetMol | Cat#:T1995 |
| NMS873 | ApexBIO | Cat#: B2168 |
| MG132 | ApexBIO | Cat#: A2585 |
| Critical Commercial Assays | | |

| | | |
|---|---|---|
| BD Pharmingen™ BrdU Flow Kits | BD biosciences | Cat#: 557892 |
| High-Capacity RNA-to-cDNA™ Kit | Thermo Fisher | Cat#: 4387406 |
| Anti-APC MicroBeads | Miltenyi biotec | Cat#: 130-097-143 |
| SuperPAGE™ 4-12% Bis-Tris Protein Gels | Epizyme | Cat#: NP0321BOX |
| Immobilon® -P PVDF Membrane | Millipore | Cat#: IPVH00010 |
| MS Columns | Miltenyi biotec | Cat#: 130-042-201 |
| Cell Strainer | BKMAMLAB. | Cat#: 110426001 |
| anti-Flag magnetic beads | Thermo Fisher | Cat#: A36798 |
| SuperSignal™ West Femto Maximum Sensitivity Substrate | Thermo Fisher | Cat#: 34095 |
| Experimental Models: Organisms/Strains | | |
| Mouse:Sel1l$^{fl/fl}$ | Sun S, et al. Proc Natl Acad Sci U S A. 2014;111:E582–E591. | PMID:24453213 |
| Mouse:JAK2$^{V617F/+}$ | Mullally A, et al. Cancer Cell. 2010 Jun 15;17(6):584-96. | PMID: 20541703 |
| Vav1-Cre | de Boer J, et al. Eur J Immunol. 2003;33:314–25. | PMID: 12548562 |
| Mouse:B6.Cg-Tg(Mx1-cre)1Cgn/J | Jackson Laboratory | Cat#:003556 |
| Mouse:SOCS2$^{3Xmcherry/+}$ | Cyagen | This study |
| Mouse:SOCS2$^{-/-}$ | Shanghai model organism | Cat#:NM-KO-190148 |
| Mouse:Rosa26$^{Zsgreen/+}$ | Cyagen | Cat#:C001262 |
| Mouse:Rosa26$^{ERADi/+}$ | Cyagen | This study |
| Mouse:Hrd1$^{fl/fl}$ | GemPharmatech | Cat#:T009546 |
| Software and Algorithms | | |
| FLOWJO | FLOWJO X.07 for windows,TreeStar, Ashland Oregon USA | https://www.flowjo.com/ |
| ImageJ | ImageJ software version 1.51j8, National Institutes of Health, Bethesda, MD, USA | https://imagej.net/ij/ |
| GraphPad Prism | GraphPad Software version 7.0 for Windows, GraphPad Software, La Jolla California USA | https://www.graphpad.com/ |

Table S2. Primers

Primers for genotyping

| Target | Primers | Seq (5'-3) |
|---|---|---|
| *Mx1-cre* | *Forward* | GCGGTCTGGCAGTAAAAACTATC |
| | *Reverse* | GTGAAACAGCATTGCTGTCACTT |
| *Rosa26* $^{ERADi/+}$ | *Forward* | AGATCTGCA GCTA TTCCTGC |
| | *Reverse* | AGGTTGATGGCCTGCTTGCCCT |
| *Hrd1* $^{fl/fl}$ | *Forward* | ATGGAGACCAGAGATCGTGTCCTACTCT |
| | *Reverse* | AAGACTACTGGAGAGTTGAGAGAAGCG |
| *Sel1L* $^{fl/fl}$ | *Forward* | CTGACTGAGGAAGGGTCTC |
| | *Reverse* | GCTAAAAACATTACAAAGGGGCA |
| *JAK2* $^{V617F/+}$ | *Forward* | CGTGCATAGTGTCTGTGGAAGTC |
| | *Reverse* | CGTGGAGAGTCTGTAAGGCTCAA |
| *SOCS2* $^{3Xmcherry/+}$ | *Forward* | GCCTCTAGTGAGAGAATGTACCCA |
| | *Reverse* | ATCTTGAGCAGCCATAGGAAAGA |
| *SOCS2* $^{-/-}$ | *Forward* | CCTCCTCACCCCTCTACTCC |
| | *Reverse* | GCTGCCTTGAGATGGAGGAC |
| *Vav-cre* | *Forward* | CAGGTTTTGGTGCACAGTCA |
| | *Reverse* | GGTGTTGTAGTTGTCCCCACT |

Primers for RT-qPCR

| Target | Primers | Seq (5'-3) |
|---|---|---|
| *mHoxb5* | *Forward* | GCTTCACATCAGCCACGATA |
| | *Reverse* | CAGGTAGCGATTGAAGTGGAAT |
| *mNeo1* | *Forward* | GCATAACCTCGGACCACAAT |
| | *Reverse* | GCTGCTCTCACAGTCAATGG |
| *mSocs2* | *Forward* | CTGCGCGAGCTCAGTCAAA |
| | *Reverse* | CAATCCGCAGGTTAGTCGGT |
| *mHrd1* | *Forward* | AGCTACTTCAGTGAACCCCACT |
| | *Reverse* | CTCCTCTACAATGCCCACTGAC |
| *mBIP* | *Forward* | CGACTTGGGGACCACCTATT |
| | *Reverse* | TTGGACGTGAGTTGGTTCTT |
| *mSel1L* | *Forward* | CTGACTGAGGAAGGGTCTC |
| | *Reverse* | CCTTTGTTCCGGTTACTTCTTG |
| *mxbp1 total* | *Forward* | AAACAGAGTAGCAGCGCAGAC |
| | *Reverse* | CAGGATCCAGCGTGTCCAT |
| *mxbp1s* | *Forward* | ACACGCTTGGGAATGGAAC |
| | *Reverse* | CCATGGGAAGATGTTCTGGG |
| *mHerp* | *Forward* | GCCGGACAACTCTAATCAGAC |
| | *Reverse* | CCCATACGTTGTGTAGCCAGA |
| *mVcp* | *Forward* | CTGCTGACCGAGTCATCAA |
| | *Reverse* | TTAGGATGGCAACACGGGA |
| *mβ-actin* | *Forward* | GGCTGTATTCCCCTCCATCG |
| | *Reverse* | CCAGTTGGTAACAATGCCATGT |
| *hSocs2* | *Forward* | GAGCCGGAGAGTCTGGTTTC |
| | *Reverse* | ATCCTGGAGGACGGATGACA |
| *hVcp* | *Forward* | ATCGGTTAATTGTTGATGAA |
| | *Reverse* | GTCTCTTCTTTCCTTTCAG |
| *hβ-actin* | *Forward* | AGAGGCATCCTCACCCTGAA |
| | *Reverse* | CACACGCAGCTCATTGTAGAA |

Table S3 Staining cocktails

2-5 ml FACS buffer was used to wash samples after each staining
HSC cocktail for basic staining and chimerism
1st staining, on ice for 30 min

| Antigen | Fluorophore | Dilution |
|---|---|---|
| Gr-1 | Biotin | 800 |
| B220 | Biotin | 400 |
| CD2 | Biotin | 400 |
| CD3 | Biotin | 400 |
| CD5 | Biotin | 400 |
| CD8 | Biotin | 400 |
| Ter-119 | Biotin | 400 |
| Sca-1 | PerCP-Cy5.5 | 200 |
| c-Kit | APC | 200 |
| CD48 | APC/Cyanine7 | 200 |
| CD150 | PE-Cy7 | 200 |
| DAPI | | 1600 |

2nd staining, on ice for 30 min

| Antigen | Fluorophore | Dilution |
|---|---|---|
| Brilliant Violet 510™ Streptavidin | N/A | 400 |

Myeloid, B/T Lymphoid lineage cocktail for staining and chimerism
1st staining, on ice for 30 min

| Antigen | Fluorophore | Dilution |
|---|---|---|
| Mac-1 | APC | 800 |
| B220 | PerCP-Cy5.5 | 200 |
| CD3 | PE-CY7 | 200 |
| Ter119 | otin-Brilliant Violet 510™ | 400 |
| DAPI | | 1600 |

2nd staining, on ice for 30 min

| Antigen | Fluorophore | Dilution |
|---|---|---|
| Brilliant Violet 510™ Streptavidin | N/A | 400 |

HSC cocktail for sorting(whole bone marrow including spine was used for sort
1st staining, on ice for 30 min

| Antigen | Fluorophore | Dilution |
|---|---|---|
| Gr-1 | Biotin | 800 |
| B220 | Biotin | 400 |
| CD2 | Biotin | 400 |
| CD3 | Biotin | 400 |
| CD5 | Biotin | 400 |
| CD8 | Biotin | 400 |
| Ter-119 | Biotin | 400 |
| Sca-1 | PerCP-Cy5.5 | 200 |
| c-Kit | APC | 200 |
| CD48 | Alexa 700 | 200 |

| | | |
|---|---|---|
| CD150 | PE-Cy7 | 200 |
| DAPI | | 1600 |

2nd staining, on ice for 30 min

| Antigen | Fluorophore | Dilution |
|---|---|---|
| Brilliant Violet 510™ Streptavidin | N/A | 400 |

3nd staining,c-Kit erichment, on ice for 30 min

| Antigen | Fluorophore | Dilution |
|---|---|---|
| Anti-APC microbeads | N/A | 7 |

Rosa26$^{ERADi/+}$ mice Toxicity HSC chimerism

1st staining, on ice for 30 min

| | | |
|---|---|---|
| Gr-1 | Biotin | 800 |
| B220 | Biotin | 400 |
| CD2 | Biotin | 400 |
| CD3 | Biotin | 400 |
| CD5 | Biotin | 400 |
| CD8 | Biotin | 400 |
| Ter-119 | Biotin | 400 |
| Sca-1 | PerCP-Cy5.5 | 200 |
| c-Kit | APC | 200 |
| CD48 | Alexa 700 | 200 |
| CD150 | PE-Cy7 | 200 |
| CD45.2 | APC/Cyanine7 | 200 |
| DAPI | | 1600 |

2nd staining, on ice for 30 min

| Antigen | Fluorophore | Dilution |
|---|---|---|
| Brilliant Violet 510™ Streptavidin | N/A | 400 |

Rosa26$^{ERADi/+}$ mice Toxicity Lineage chimerism

1st staining, on ice for 30 min

| Antigen | Fluorophore | Dilution |
|---|---|---|
| Mac-1 | APC | 800 |
| B220 | PerCP-Cy5.5 | 200 |
| CD3 | PE-CY7 | 200 |
| CD45.2 | APC/Cyanine7 | 200 |
| DAPI | | 1600 |

2nd staining, on ice for 30 min

| Antigen | Fluorophore | Dilution |
|---|---|---|
| Brilliant Violet 510™ Streptavidin | N/A | 400 |

Rosa26$^{ERADi/+}$ mice Lineage Bleeding

| Antigen | Fluorophore | Dilution |
|---|---|---|

| | | |
|---|---|---|
| Mac-1 | APC | 800 |
| B220 | PerCP-Cy5.5 | 200 |
| CD3 | PE-CY7 | 200 |
| CD45.1 | Brilliant Violet 510™ | 200 |
| CD45.2 | APC/Cyanine7 | 200 |
| DAPI | | 400 |

Rosa26$^{ERADi/+}$ mice HSC Staining for BRDU, on ice for 30 min
1st staining, on ice for 30 min

| Antigen | Fluorophore | Dilution |
|---|---|---|
| Gr-1 | Biotin | 600 |
| B220 | Biotin | 200 |
| CD2 | Biotin | 200 |
| CD3 | Biotin | 400 |
| CD5 | Biotin | 200 |
| CD8 | Biotin | 200 |
| Ter-119 | Biotin | 200 |
| Sca-1 | PerCP-Cy5.5 | 200 |
| c-Kit | PECY7 | 200 |
| CD48 | Alexa700 | 200 |
| CD150 | BV711 | 100 |
| DAPI | | 1600 |

2nd staining, on ice for 30 min

| Antigen | Fluorophore | Dilution |
|---|---|---|
| Brilliant Violet 510™ Streptavidin | N/A | 400 |

CD200 or CD86 HSC cocktail for basic staining and sorting
1st staining, on ice for 30 min

| Antigen | Fluorophore | Dilution |
|---|---|---|
| Gr-1 | FITC | 800 |
| B220 | FITC | 400 |
| CD2 | FITC | 400 |
| CD3 | FITC | 400 |
| CD5 | FITC | 400 |
| CD8 | FITC | 400 |
| Ter-119 | FITC | 400 |
| Sca-1 | PerCP-Cy5.5 | 200 |
| c-Kit | APC | 200 |
| CD48 | APC/Cyanine7 | 200 |
| CD150 | PE | 200 |
| CD200 or CD86 | PE-CY7 | 200 |
| DAPI | | 1600 |

2nd staining,c-Kit erichment, on ice for 30 min

| Antigen | Fluorophore | Dilution |
|---|---|---|
| Anti-APC microbeads | N/A | 7 |

CD200 Lineage Bleeding

| Antigen | Fluorophore | Dilution |
|---|---|---|

| | | |
|---|---|---|
| Mac-1 | APC | 800 |
| Gr-1 | FITC | 800 |
| B220 | PerCP-Cy5.5 | 200 |
| CD3 | PE | 200 |
| CD200 or CD86 | PE-CY7 | 200 |
| CD45.1 | Brilliant Violet 510™ | 200 |
| CD45.2 | APC/Cyanine7 | 200 |
| DAPI | | 400 |

CD200  HSC chimerism

| Antigen | Fluorophore | Dilution |
|---|---|---|
| Gr-1 | FITC | 800 |
| B220 | FITC | 400 |
| CD2 | FITC | 400 |
| CD3 | FITC | 400 |
| CD5 | FITC | 400 |
| CD8 | FITC | 400 |
| Ter-119 | FITC | 400 |
| Sca-1 | PerCP-Cy5.5 | 200 |
| c-Kit | APC | 200 |
| CD48 | APC/Cyanine7 | 200 |
| CD150 | PE | 200 |
| CD45.1 | BV-510 | 200 |
| CD200 | PE-CY7 | 200 |

CD200 Lineage chimerism

| Antigen | Fluorophore | Dilution |
|---|---|---|
| Mac-1 | APC | 800 |
| Gr-1 | FITC | 800 |
| B220 | PerCP-Cy5.5 | 200 |
| CD3 | PE | 200 |
| CD200 | PE-CY7 | 200 |
| CD45.1 | Brilliant Violet 510™ | 200 |
| CD45.2 | APC/Cyanine7 | 200 |
| DAPI | | 400 |

Rosa26$^{ERADi/+}$ mice CD200 or CD86 HSC cocktail for basic staining
1st staining, on ice for 30 min

| Antigen | Fluorophore | Dilution |
|---|---|---|
| Gr-1 | Biotin | 800 |
| B220 | Biotin | 400 |
| CD2 | Biotin | 400 |
| CD3 | Biotin | 400 |
| CD5 | Biotin | 400 |
| CD8 | Biotin | 400 |
| Ter-119 | Biotin | 400 |
| Sca-1 | PerCP-Cy5.5 | 200 |
| c-Kit | APC | 200 |
| CD48 | Alexa 700 | 200 |
| CD150 | BV711 | 200 |
| CD200 or CD86 | PE-CY7 | 200 |

| | | |
|---|---|---|
| DAPI | | 1600 |

2nd staining, on ice for 30 min

| Antigen | Fluorophore | Dilution |
|---|---|---|
| Brilliant Violet 510™ Streptavidin | N/A | 400 |

SOCS2 3XmCherry mice cocktail for HSC chimerism staining

| Antigen | Fluorophore | Dilution |
|---|---|---|
| Gr-1 | FITC | 800 |
| B220 | FITC | 400 |
| CD2 | FITC | 400 |
| CD3 | FITC | 400 |
| CD5 | FITC | 400 |
| CD8 | FITC | 400 |
| Ter-119 | FITC | 400 |
| Sca-1 | PerCP-Cy5.5 | 200 |
| c-Kit | APC | 200 |
| CD48 | Alexa 700 | 200 |
| CD150 | PE-CY7 | 200 |
| CD45.1 | Brilliant Violet 510™ | 200 |
| CD45.2 | APC/Cyanine7 | 200 |
| DAPI | | 1600 |

SOCS2 3XmCherry mice cocktail for lineage chimerism staining

| Antigen | Fluorophore | Dilution |
|---|---|---|
| Mac-1 | APC | 800 |
| B220 | PerCP-Cy5.5 | 200 |
| CD3 | PE-CY7 | 200 |
| CD45.1 | Brilliant Violet 510™ | 200 |
| CD45.2 | APC/Cyanine7 | 200 |
| DAPI | | 1600 |

SOCS2 3XmCherry mice Lineage Bleeding

| Antigen | Fluorophore | Dilution |
|---|---|---|
| Mac-1 | APC | 800 |
| B220 | PerCP-Cy5.5 | 200 |
| CD3 | PE-CY7 | 200 |
| CD45.1 | Brilliant Violet 510™ | 200 |
| CD45.2 | APC/Cyanine7 | 200 |
| DAPI | | 400 |

### Supplementary Figures

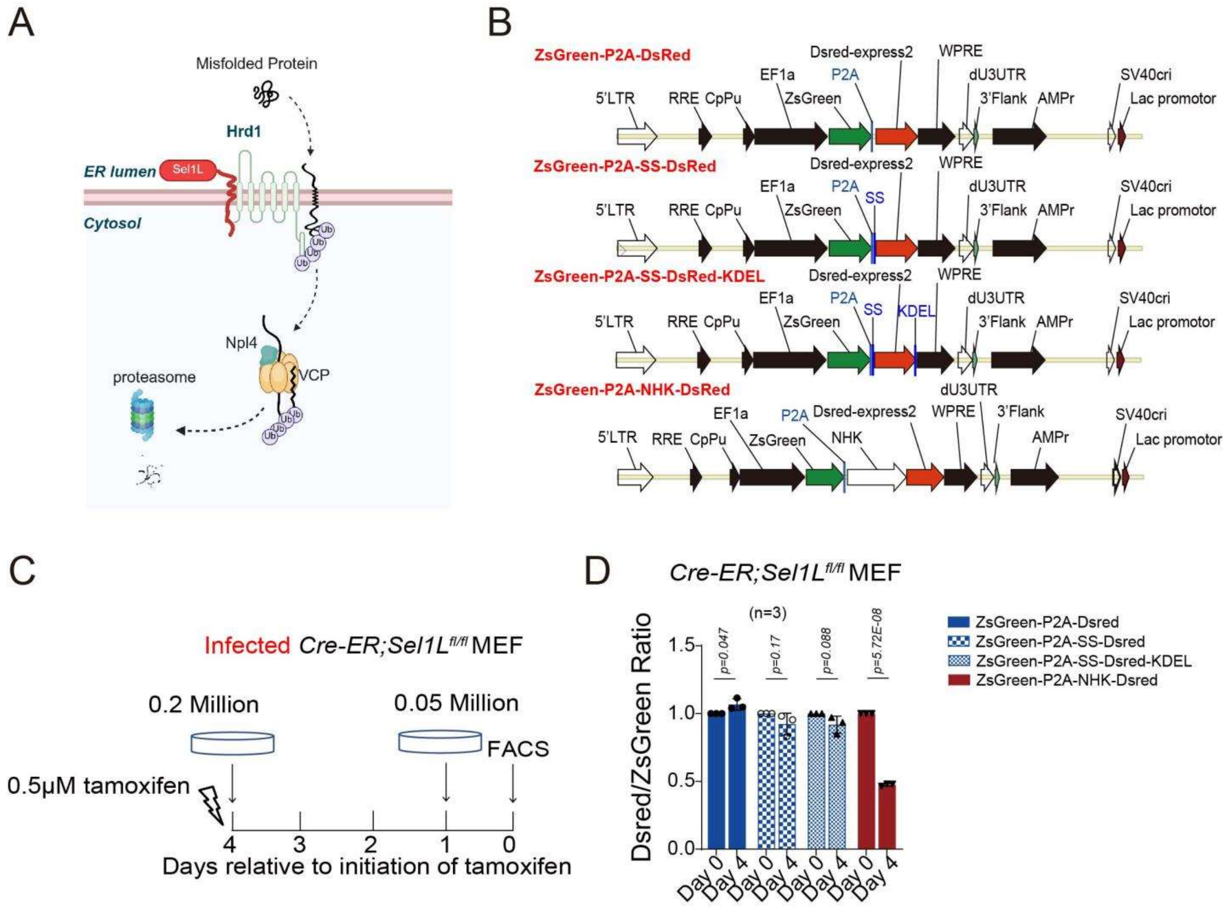

**Fig. S1. A dual-fluorescence reporter enables quantitative, real-time measurement of ERAD activity in vitro and in vivo. (A)** Schematic model of ERAD targeting misfolded proteins for degradation. **(B)** ERAD activity reporter and control constructs for cell-based assays. **(C-D)** *ER-Cre; Sel1L$^{fl/fl}$* MEFs were infected with PEZ-NHK or PEZ-control lentiviral constructs and treated with tamoxifen (2 μg/ml) for 0-4 days to induce Sel1L deletion and evaluate ERAD activity. Data represent mean±s.d. from three independent experiments. Statistical significance was determined using unpaired two-tailed Student's t-test.

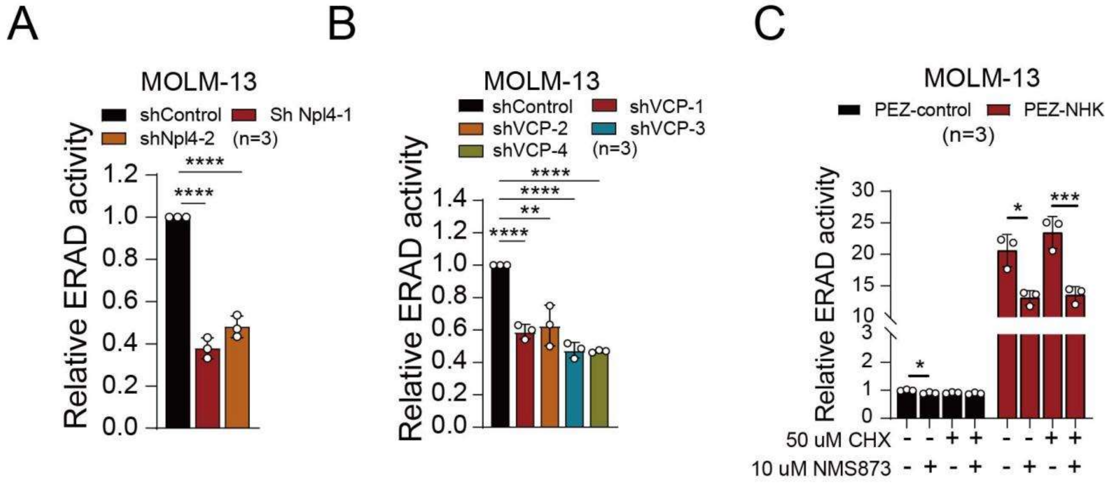

**Fig. S2. The ERAD reporter reliably detects ERAD perturbations in human hematopoietic cells.** **(A)** MOLM-13 cells were infected with shNpl4 or control lentivirus, selected with puromycin for 48 hours, and analyzed by flow cytometry to evaluate ERAD activity. **(B)** MOLM-13 cells were infected with shVCP or control lentivirus, selected with puromycin for 48 hours, and analyzed by flow cytometry to evaluate ERAD activity. **(C)** MOLM-13 cells were infected with PEZ-NHK (ERAD reporter) or PEZ-control lentivirus and treated with NMS873 (10 µM) and cycloheximide (50 µM) for 24 hours. ERAD activity was assessed by flow cytometry. Data represent mean±s.d. from three independent experiments. Statistical significance was determined using unpaired two-tailed Student's t-test.

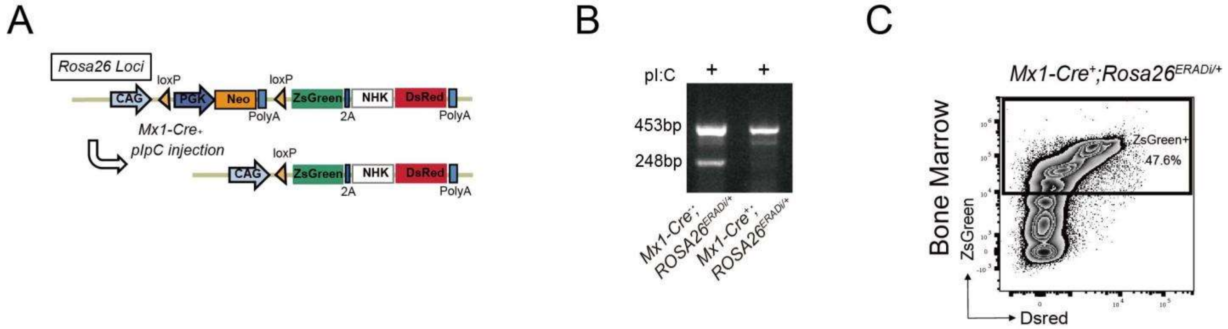

**Fig. S3. The ERAD reporter exhibits minimal toxicity and does not perturb steady-state hematopoiesis. (A)** Schematic representation of the ERAD reporter mice. **(B)** Genotyping of whole bone marrow cells from *Mx1-Cre*$^{+}$; *Rosa26*$^{ERADi/+}$ mice and control littermates after poly(I:C) induction showed deletion of the 248 bp band, indicating successful Cre-mediated recombination. **(C)** Whole bone marrow cells from *Mx1-Cre*$^{+}$; *Rosa26*$^{ERADi/+}$ mice were analyzed for ZsGreen and DsRed expression by flow cytometry following poly(I:C) induction.

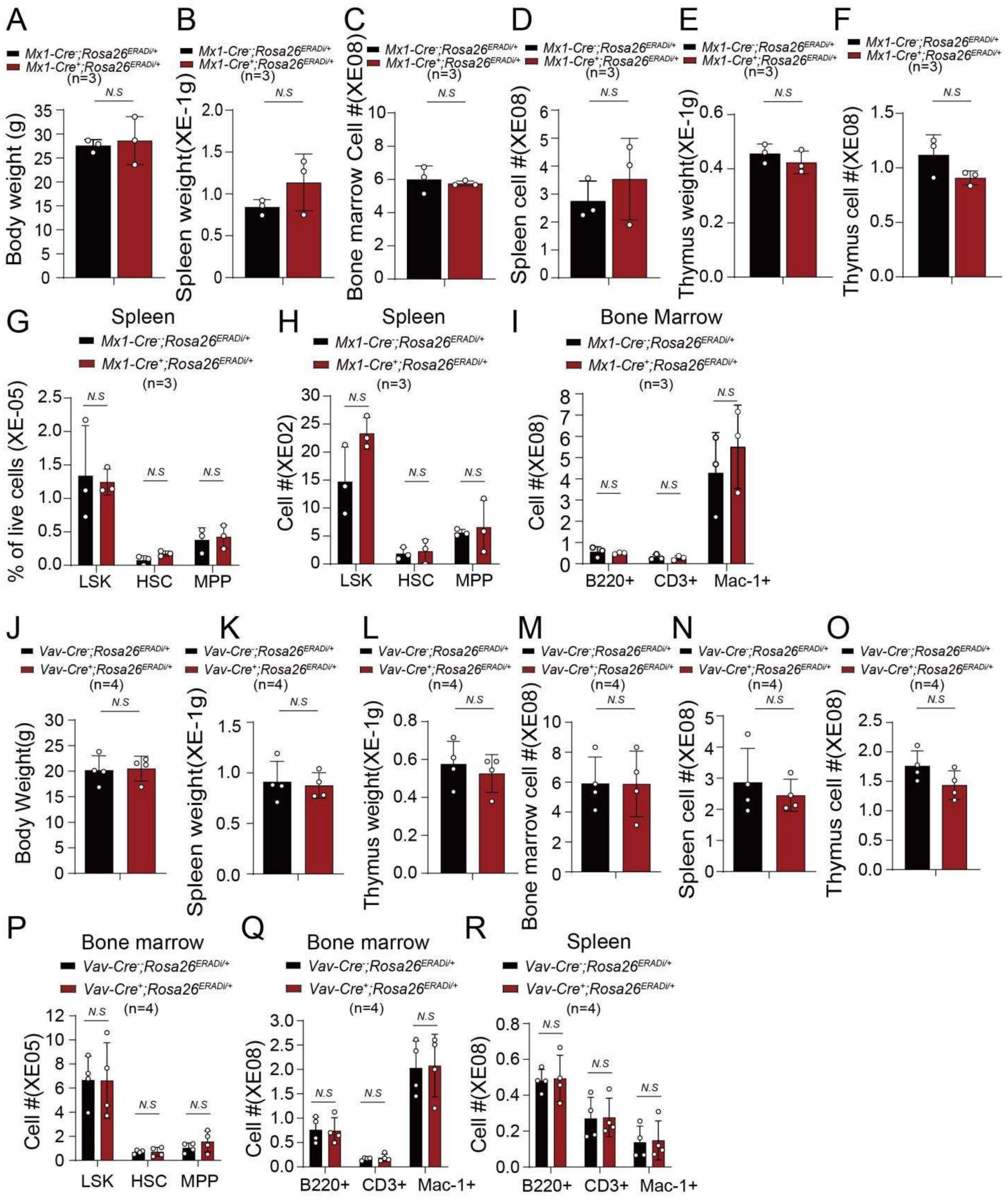

**Fig. S4. The ERAD reporter exhibits minimal toxicity and does not perturb steady-state hematopoiesis.** 6-8 weeks old *Mx1-Cre*$^{-}$*; Rosa26*$^{ERADi/+}$ and *Mx1-Cre*$^{+}$*; Rosa26*$^{ERADi/+}$ mice were injected with poly(I:C) every other day for a total of three doses. Two weeks post-injection, body weight **(A)**, spleen weight **(B)**, cellularity of bone marrow **(C)** and spleen **(D)**, thymus weight **(E)** and cellularity **(F)**, frequency and numbers of hematopoietic stem and progenitor cells (HSPCs) in the spleen **(G-H)**, and numbers of mature blood cells in the bone marrow **(I)** were assessed. 6 to 8 weeks old *Vav1-Cre*$^{+}$*; Rosa26*$^{ERADi/+}$ and *Vav1-Cre*$^{-}$*; Rosa26*$^{ERADi/+}$ mice were analyzed for body weight **(J)**, spleen **(K)** and thymus **(L)** weight, cellularity of bone marrow **(M)**, spleen **(N)** and thymus **(O)**, numbers of HSPCs **(P)**, and mature blood cells in the bone marrow and spleen **(Q-R)**. Data represent mean±s.d. from at least three independent experiments. Statistical significance was determined using unpaired two-tailed Student's t-test.

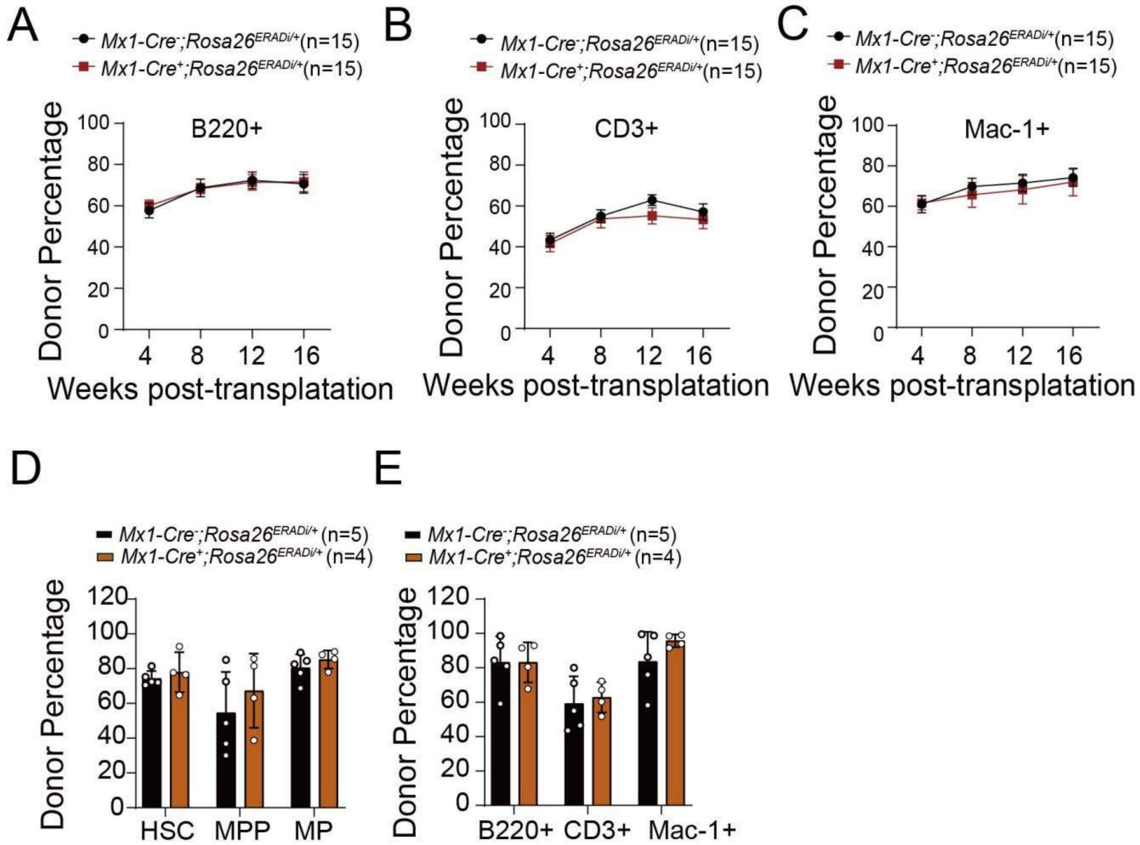

**Fig. S5. ERAD reporter expression does not impair HSC long-term repopulation capacity.** 6-8 weeks old *Mx1-Cre*$^{-}$*; Rosa26*$^{ERADi/+}$ and *Mx1-Cre*$^{+}$*; Rosa26*$^{ERADi/+}$ mice were injected with poly(I:C) every other day for a total of three doses. Two weeks post-injection, 5x$10^5$ CD45.2$^{+}$ whole bone marrow cells from *Mx1-Cre*$^{+}$*; Rosa26*$^{ERADi/+}$ and control (+/+) mice transplanted together with 5x$10^5$ CD45.1$^{+}$ wild type bone marrow cells into lethally irradiated CD45.1$^{+}$ wild-type recipients. **(A-C)** The contribution of CD45.2 cells in B (B220$^{+}$; **A**), T (CD3$^{+}$; **B**) and myeloid (Mac1$^{+}$; **C**) cells from peripheral blood was analyzed every 4 weeks for 16 weeks. **(D-E)** The contribution of CD45.2 cells in HSPC **(D)**, and mature lineage cells **(E)** in the bone marrow was analyzed at 16 weeks post-transplantation. Data represent mean±s.d. from three independent experiments. Statistical significance was determined using unpaired two-tailed Student's t-test.

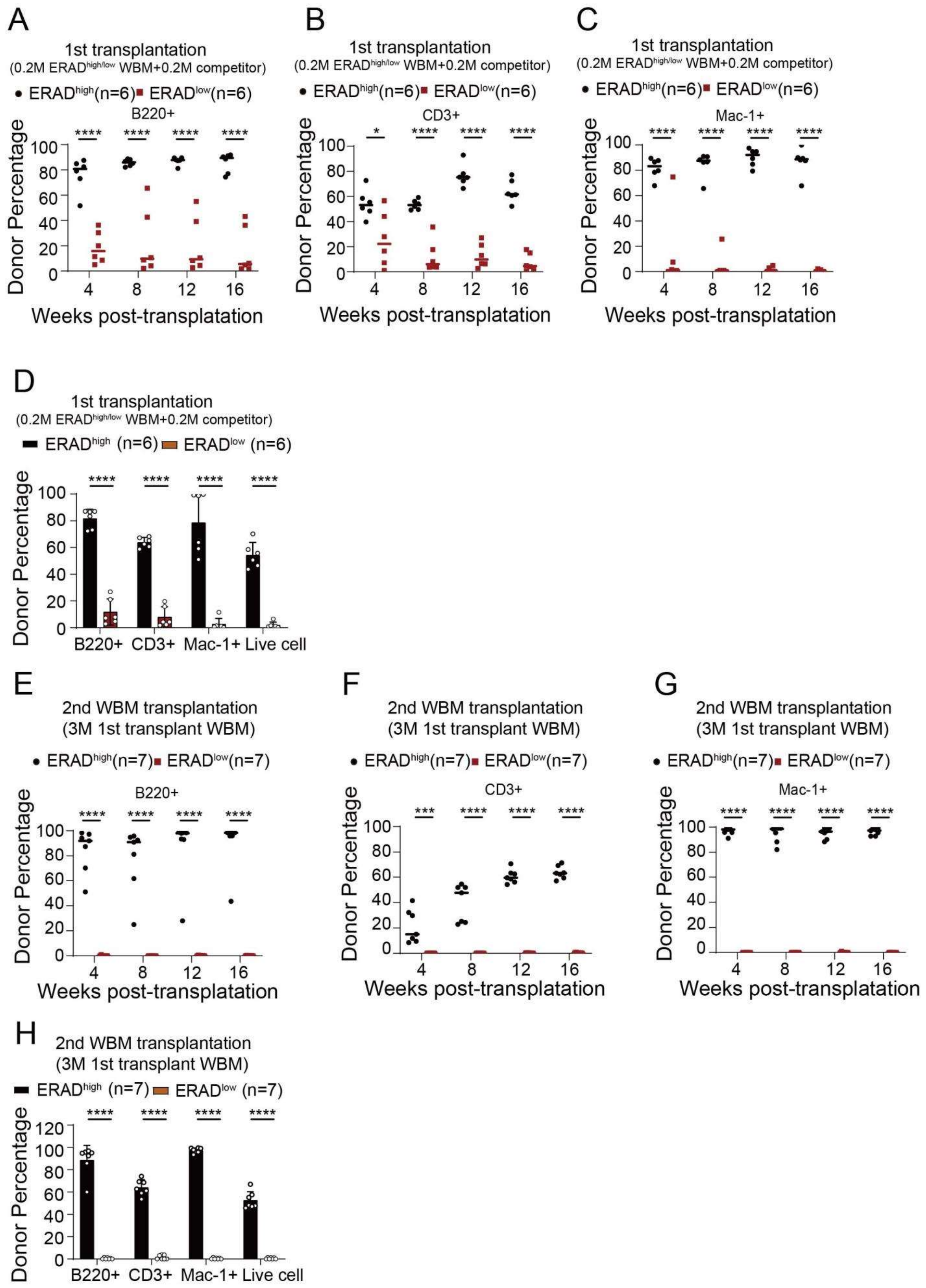
A
1st transplantation
(0.2M ERADhigh/low WBM+0.2M competitor)
• ERADhigh(n=6) ■ ERADlow(n=6)
B220+
Donor Percentage
Weeks post-transplatation
B
1st transplantation
(0.2M ERADhigh/low WBM+0.2M competitor)
• ERADhigh(n=6) ■ ERADlow(n=6)
CD3+
Donor Percentage
Weeks post-transplatation
C
1st transplantation
(0.2M ERADhigh/low WBM+0.2M competitor)
• ERADhigh(n=6) ■ ERADlow(n=6)
Mac-1+
Donor Percentage
Weeks post-transplatation
D
1st transplantation
(0.2M ERADhigh/low WBM+0.2M competitor)
ERADhigh (n=6) ERADlow (n=6)
Donor Percentage
B220+ CD3+ Mac-1+ Live cell
E
2nd WBM transplantation
(3M 1st transplant WBM)
• ERADhigh(n=7) ■ ERADlow(n=7)
B220+
Donor Percentage
Weeks post-transplatation
F
2nd WBM transplantation
(3M 1st transplant WBM)
• ERADhigh(n=7) ■ ERADlow(n=7)
CD3+
Donor Percentage
Weeks post-transplatation
G
2nd WBM transplantation
(3M 1st transplant WBM)
• ERADhigh(n=7) ■ ERADlow(n=7)
Mac-1+
Donor Percentage
Weeks post-transplatation
H
2nd WBM transplantation
(3M 1st transplant WBM)
ERADhigh (n=7) ERADlow (n=7)
Donor Percentage
B220+ CD3+ Mac-1+ Live cell

**Fig. S6. ERAD activity prospectively identifies bone marrow cells with superior reconstitution potential**. 6-8 weeks old *Mx1-Cre*$^{+}$; *Rosa26*$^{ERADi/+}$ mice were injected with poly(I:C) every other day for a total of three doses. Two weeks post-injection, whole bone marrow cells were sorted by flow cytometry into 2x$10^5$ ERAD$^{high}$ and ERAD$^{low}$ populations and transplanted together with 2x$10^5$ CD45.1$^{+}$ wild type bone marrow cells into lethally irradiated CD45.1$^{+}$ wild-type recipients. **(A-C)** The contribution of CD45.2 cells in B (B220$^{+}$; **A**), T (CD3$^{+}$; **B**) and myeloid (Mac1$^{+}$; **C**) cells from peripheral blood was analyzed every 4 weeks for 16 weeks. **(D)** The contribution of CD45.2 cells in mature lineage cells in the bone marrow was analyzed at 16 weeks post-transplantation. For secondary transplantation, 3 million whole bone marrow cells from the primary recipients transplanted with 2x$10^5$ ERAD$^{high}$ and ERAD$^{low}$ populations were transplanted into irradiated secondary receipt CD45.1 mice. **(E-G)** The contribution of CD45.2 cells in B (B220$^{+}$; **E**), T (CD3$^{+}$; **F**) and myeloid (Mac1$^{+}$; **G**) cells from peripheral blood was analyzed every 4 weeks for 16 weeks. **(H)** The contribution of CD45.2 cells in mature lineage cells in the bone marrow was analyzed at 16 weeks post-transplantation. Data represent mean±s.d. from three independent experiments. Statistical significance was determined using unpaired two-tailed Student's t-test.

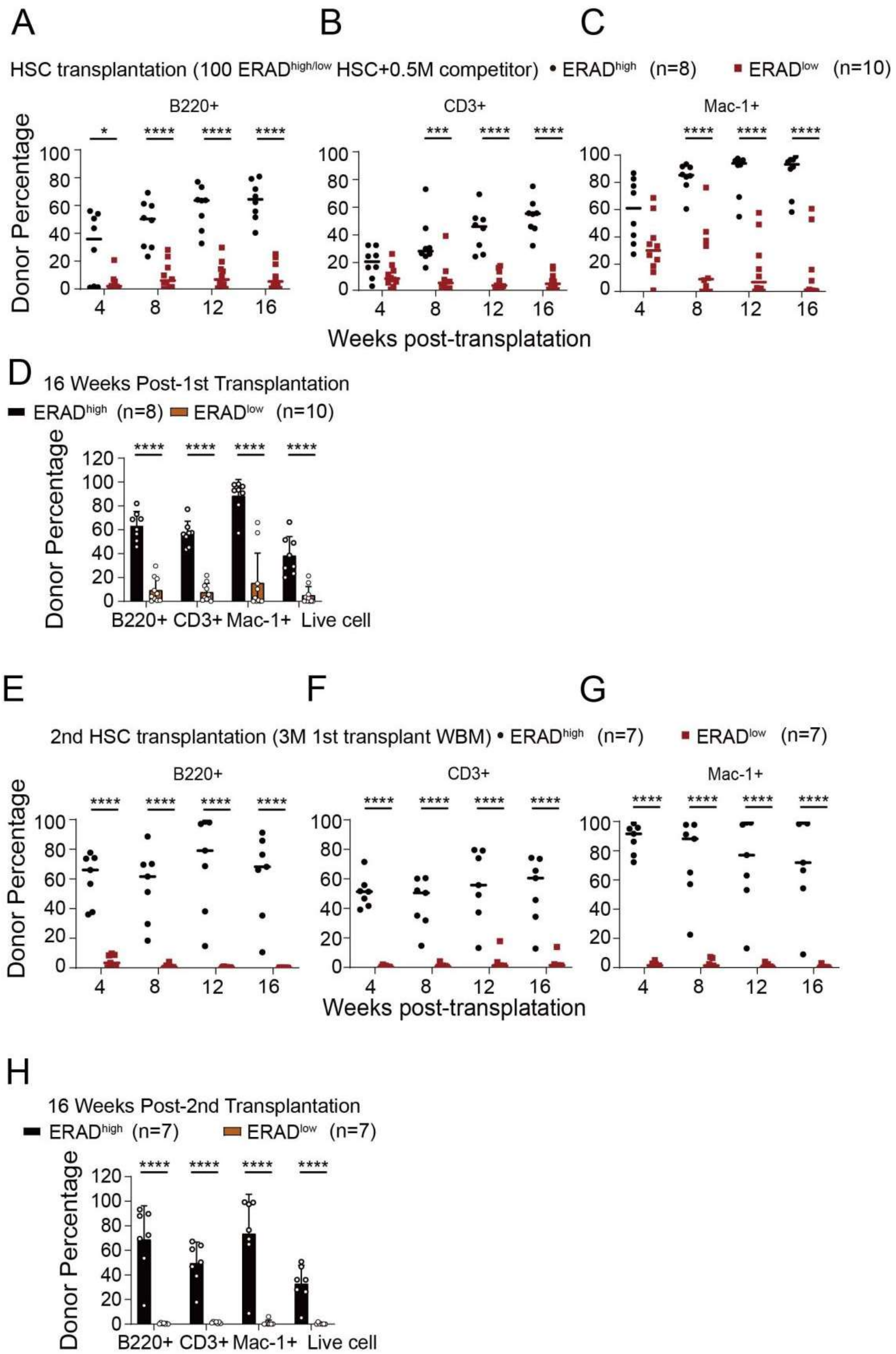

**Fig. S7. High ERAD activity marks HSCs with durable self-renewal and multilineage reconstitution capacity.** FACS-purified ERAD$^{high}$ and ERAD$^{low}$ ZsGreen$^{+}$ HSCs from *Mx1-Cre$^{+}$; Rosa26$^{ERADi/+}$* mice were transplanted into irradiated receipt mice along with 0.5 million CD45.2 whole bone marrow competitors. **(A-D)** The contribution of ZsGreen$^{+}$ cells in B (B220$^{+}$; **A**), T (CD3$^{+}$; **B**) and myeloid (Mac1$^{+}$; **C**) cells from peripheral blood was analyzed every 4 weeks for

16 weeks. **(D)** The contribution of ZsGreen$^+$ cells in HSC and progenitors in the bone marrow was analyzed at 16 weeks post-transplantation. For secondary transplantation, 3 million whole bone marrow cells from the primary recipients transplanted with 100 ERAD$^{high}$ and ERAD$^{low}$ ZsGreen$^+$ HSCs were transplanted into irradiated secondary receipt CD45.2 mice. **(E-G)** The contribution of ZsGreen$^+$ cells in B (B220$^+$; **E**), T (CD3$^+$; **F**) and myeloid (Mac1$^+$; **G**) cells from peripheral blood was analyzed every 4 weeks for 16 weeks. **(H)** The contribution of ZsGreen$^+$ cells in mature lineage cells in the bone marrow was analyzed at 16 weeks post-transplantation. Data represent mean±s.d. from three independent experiments. Statistical significance was determined using unpaired two-tailed Student's t-test.

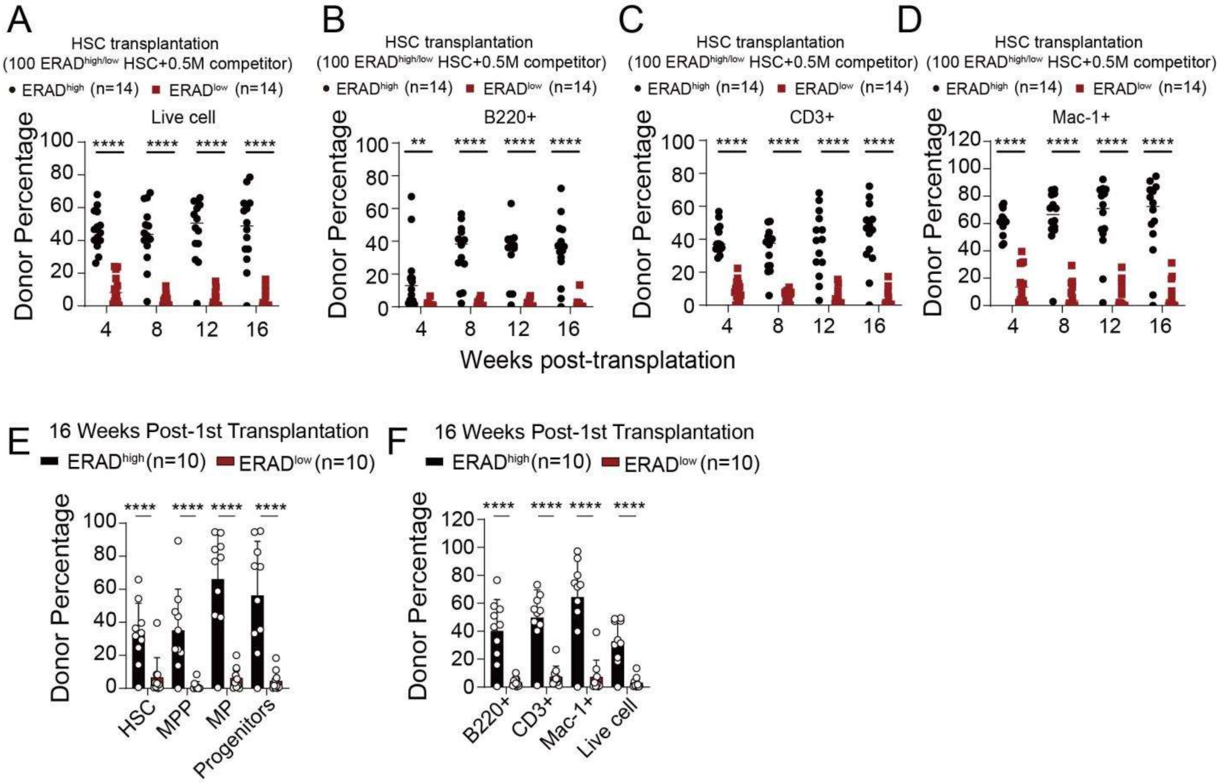

**Fig. S8. Validation of ERAD-dependent functional stratification of HSCs in the Vav1-Cre system**. FACS-purified ERAD$^{high}$ and ERAD$^{low}$ ZsGreen$^{+}$ HSCs from CD45.2 *Vav1-Cre*$^{+}$*; Rosa26*$^{ERADi/+}$ mice were transplanted into irradiated receipt mice along with 0.5 million CD45.2 whole bone marrow competitors. **(A-D)** The contribution of ZsGreen$^{+}$ cells in total CD45$^{+}$**(A)**, B (B220$^{+}$; **B**), T (CD3$^{+}$; **C**) and myeloid (Mac1$^{+}$; **D**) cells from peripheral blood was analyzed every 4 weeks for 16 weeks. **(E-F)** The contribution of ZsGreen$^{+}$ cells in HSC, progenitors **(E)** and mature lineage **(F)** cells in the bone marrow was analyzed at 16 weeks post-transplantation. Data represent mean±s.d. from three independent experiments. Statistical significance was determined using unpaired two-tailed Student's t-test.

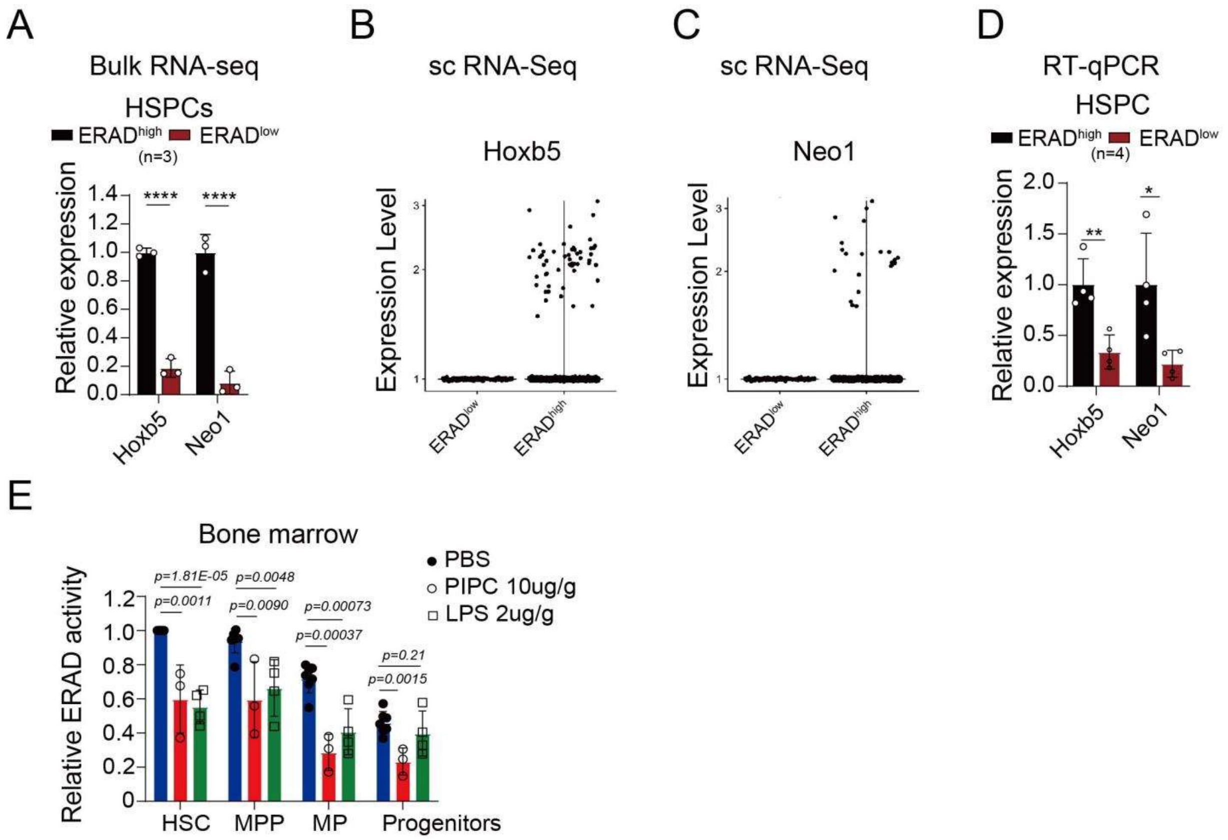

**Fig. S9. High ERAD activity correlates with quiescence-associated transcriptional signatures in HSCs.** 6-8 weeks old *Mx1-Cre*$^{+}$*; Rosa26*$^{ERADi/+}$ mice were injected with poly(I:C) every other day for a total of three doses. Two weeks post-injection, ERAD$^{high}$ and ERAD$^{low}$ HSPCs were isolated by flow cytometry for bulk RNA-sequencing. ERAD$^{high}$ and ERAD$^{low}$ lineage$^{-}$ c-Kit$^{+}$ cells were sorted for single cell RNA-sequencing. For each sample, bone marrow cells from 1-2 mice were pooled and sorted into ERAD$^{high}$ and ERAD$^{low}$ samples; *n* refers to the number of biological pools. **(A)** mRNA expression of *Hoxb5* and *Neo1* in ERAD$^{high}$ and ERAD$^{low}$ HSPCs measured by bulk RNA-seq. **(B-C)** mRNA expression of *Hoxb5* **(B)** and *Neo1* **(C)** in ERAD$^{high}$ and ERAD$^{low}$ lineage$^{-}$ c-Kit$^{+}$ cells measured by single cell RNA-seq. **(D)** Validation of expression level of *Hoxb5* and *Neo1* in ERAD$^{high}$ and ERAD$^{low}$ HSPCs by qRT-PCR. **(E)** Mice were treated with PIPC (10 μg/g, i.p.) for 48 h or LPS (2 μg/g, i.p.) for 24 h. Relative ERAD activity was analyzed by flow cytometry in HSC, MPP, MP, and LK populations. Data represent mean±s.d. from three independent experiments. Statistical significance was determined using unpaired two-tailed Student's t-test.

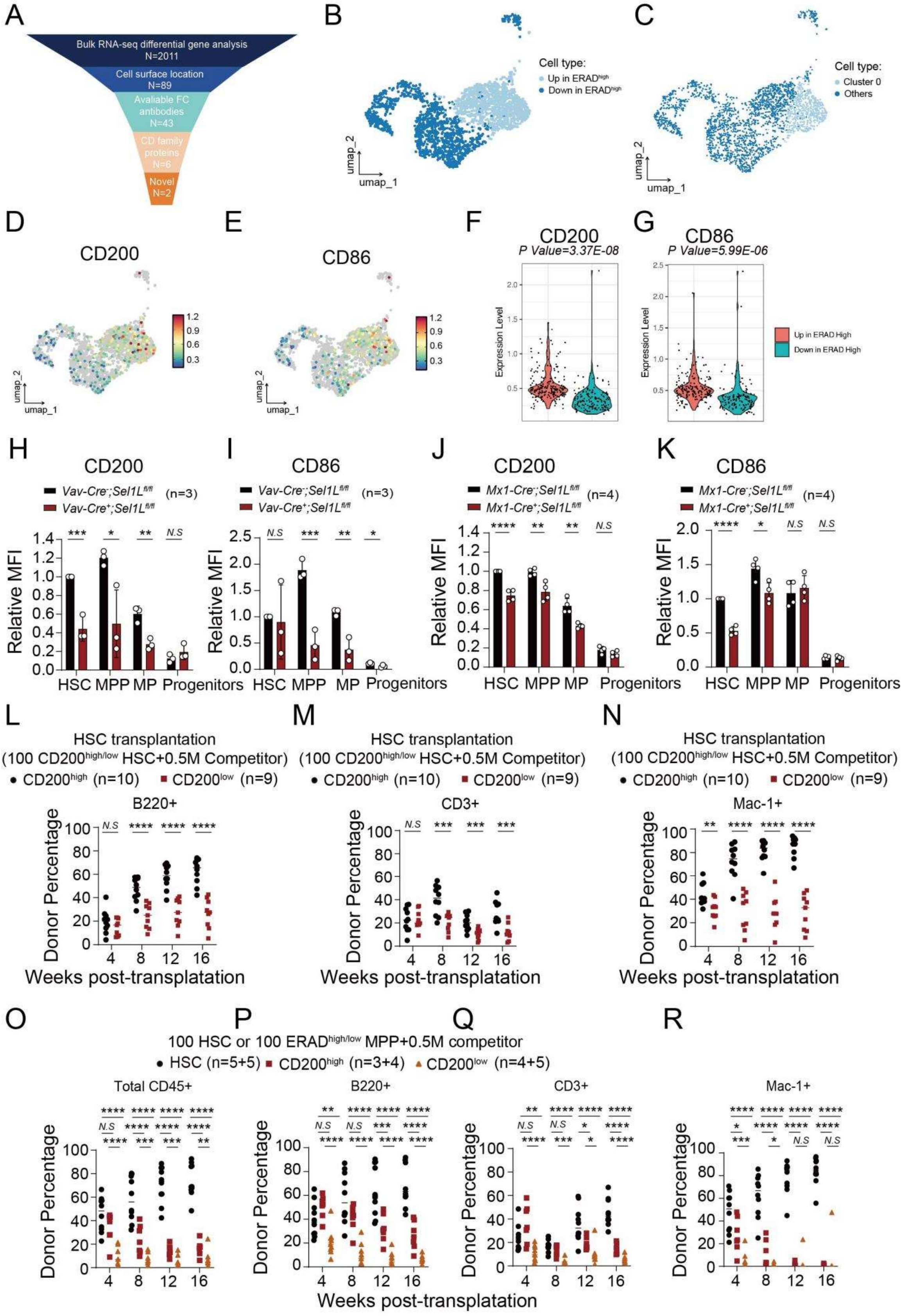
A
Bulk RNA-seq differential gene analysis N=2011
Cell surface location N=89
Avaliable FC antibodies N=43
CD family proteins N=6
Novel N=2
B
Cell type:
Up in ERADhigh
Down in ERADhigh
umap_1
umap_2
C
Cell type:
Cluster 0
Others
D
CD200
E
CD86
F
CD200
P Value=3.37E-08
Expression Level
G
CD86
P Value=5.99E-06
Up in ERAD High
Down in ERAD High
H
CD200
Vav-Cre-;Sel1Lfl/fl
Vav-Cre+;Sel1Lfl/fl
(n=3)
Relative MFI
HSC MPP MP Progenitors
I
CD86
J
CD200
Mx1-Cre-;Sel1Lfl/fl
Mx1-Cre+;Sel1Lfl/fl
(n=4)
K
CD86
L
M
N
HSC transplantation
(100 CD200high/low HSC+0.5M Competitor)
CD200high (n=10) CD200low (n=9)
B220+
CD3+
Mac-1+
Donor Percentage
Weeks post-transplatation
O
P
Q
R
100 HSC or 100 ERADhigh/low MPP+0.5M competitor
HSC (n=5+5) CD200high (n=3+4) CD200low (n=4+5)
Total CD45+
B220+
CD3+
Mac-1+
Donor Percentage
Weeks post-transplatation

**Fig. S10. CD200 faithfully mirrors ERAD activity dynamics and distinguishes functional HSC subsets. (A)** Screening strategy of novol endogenous cell surfce markers. **(B)** Uniform manifold approximation and projection (UMAP) plot of “Up in ERAD$^{high}$” and “Down in ERAD$^{high}$” cell types in the HSCs. Clusters 0, 1, and 5 are classified into the cell population “Up in ERAD$^{high}$” while clusters 2-7 are classified into the cell population “Down in ERAD$^{high}$”. **(C)** UMAP plot of “Cluster 0” and “Others” cell types in the HSCs. Cluster 0 is classified separately, while all other clusters (2, 3, 4, 5, 6, and 7) are grouped into the “Others” category. **(D-E)** The distribution of cells with different Cd200 **(D)** expression intensities or Cd86 **(E)** expression intensities on the UMAP plot. **(F - G)** Comparison of Cd200 **(F)** and Cd86 **(G)** expression levels in “Up in ERAD$^{high}$”and “Down in ERAD$^{high}$” populations. **(H-I)** Flow cytometric analysis of CD200 **(H)** and CD86 **(I)** expression in HSPCs from *Vav1-Cre*$^{+}$*; Sel1L*$^{fl/fl}$ (Sel1L KO) and control (+/+) mice. **(J-K)** Flow cytometric analysis of CD200 **(J)** and CD86 **(K)** expression in HSPCs from *Mx1-Cre*$^{+}$*; Sel1L* $^{fl/fl}$ (Sel1L KO) and control (+/+) mice. FACS-purified CD200$^{high}$ and CD200$^{low}$ HSCs from CD45.2 mice were transplanted into irradiated receipt mice along with 0.5 million CD45.1 whole bone marrow competitors. **(L-N)** The contribution of CD45.2 cells in B (B220$^{+}$; **L**), T (CD3$^{+}$; **M**) and myeloid (Mac1$^{+}$; **N**) cells from peripheral blood was analyzed every 4 weeks for 16 weeks. FACS-purified HSCs, CD200$^{high}$ and CD200$^{low}$ MPPs from CD45.2 mice were transplanted into irradiated receipt mice along with 0.5 million CD45.1 whole bone marrow competitors. **(O-R)** The contribution of CD45.2 cells in total CD45$^{+}$ **(O)**, B (B220$^{+}$; **P**), T (CD3$^{+}$; **Q**) and myeloid (Mac1$^{+}$; **R**) cells from peripheral blood was analyzed every 4 weeks for 16 weeks. Data represent mean±s.d. from three independent experiments. Statistical significance was determined using unpaired two-tailed Student’s t-test.

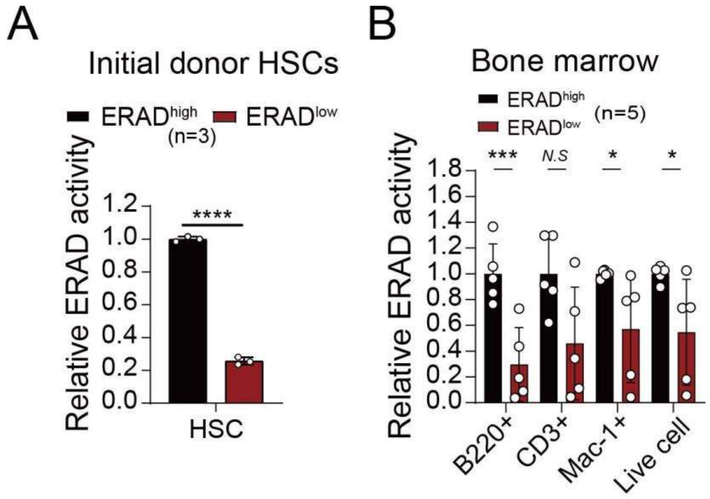

**Fig. S11. ERAD activity is a heritable proteostatic characteristic maintained through HSC self-renewal.** FACS-purified ERAD$^{high}$ and ERAD$^{low}$ ZsGreen$^{+}$ HSCs from *Mx1-Cre*$^{+}$*;* *Rosa26*$^{ERADi/+}$ mice were transplanted into irradiated receipt mice along with 0.5 million CD45.2 whole bone marrow competitors. **(A)** Comparison of relative ERAD activity between initial donor ERAD$^{high}$ and ERAD$^{low}$ HSCs. **(B)** The relative ERAD activity in donor-derived mature lineage cells in the bone marrow was analyzed 8 weeks post-transplantation. Data represent mean±s.d. from three independent experiments. Statistical significance was determined using unpaired two-tailed Student's t-test.

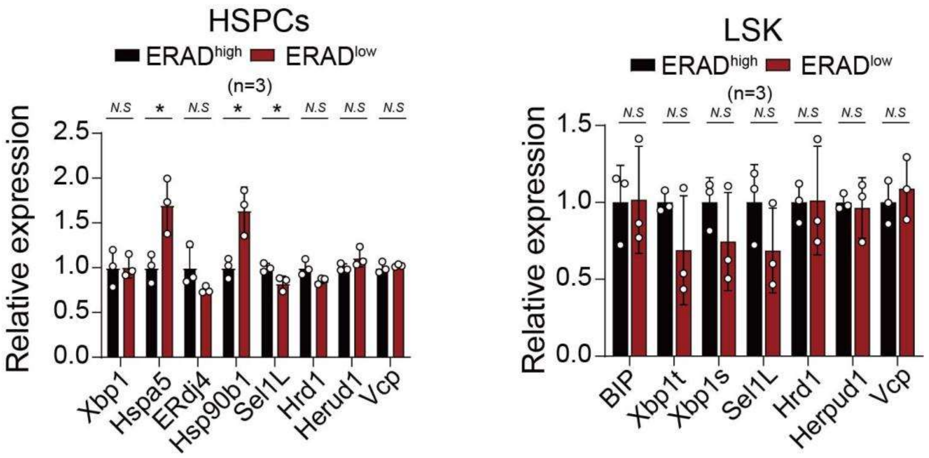

**Fig. S12. ERAD activity heterogeneity cannot be explained by canonical UPR gene expression.** 6-8 weeks old *Mx1-Cre*$^{+}$*; Rosa26*$^{ERADi/+}$ mice were injected with poly(I:C) every other day for a total of three doses. Two weeks post-injection, ERAD$^{high}$ and ERAD$^{low}$ HSPCs were isolated by flow cytometry. For each sample, bone marrow cells from 1-2 mice were pooled and sorted into ERAD$^{high}$ and ERAD$^{low}$ HSPCs; *n* refers to the number of biological pools. **(A-B)** mRNA expression of genes involved in the UPR and ERAD pathways in ERAD$^{high}$ and ERAD$^{low}$ HSPCs, as measured by RNA-seq **(A)** and validated by qRT-PCR **(B)**. Data represent mean±s.d. from three independent experiments. Statistical significance was determined using unpaired two-tailed Student's t-test.

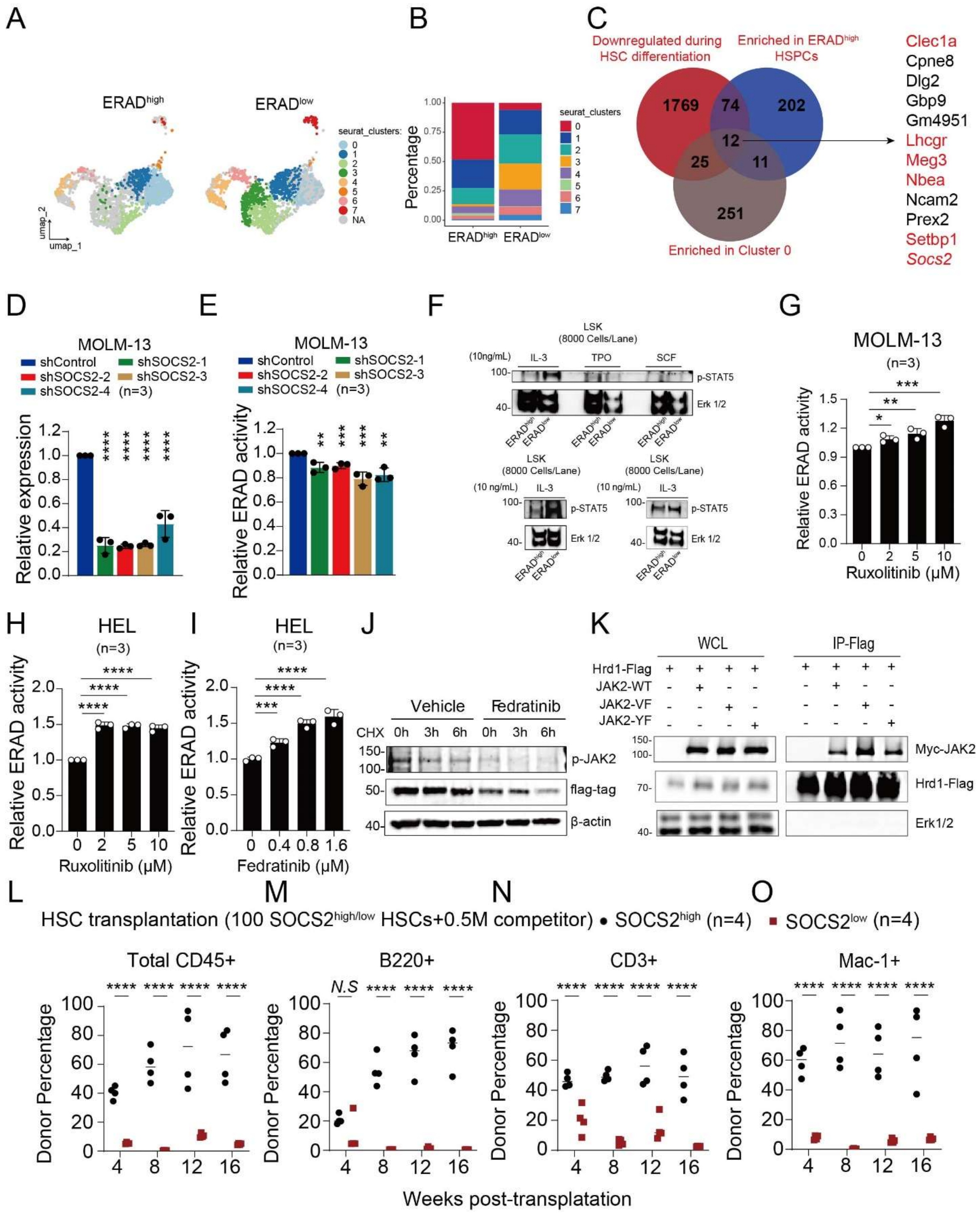

**Fig. S13. SOCS2 regulates ERAD activity heterogeneity via JAK2 signaling and predicts HSC functional potential**. **(A)** UMAP plot of the eight clusters identified in the HSCs. **(B)** Proportional differences for each cluster of $ERAD^{high}$ and $ERAD^{low}$ HSCs. **(C)** Venn diagram and 12 top overlapping genes (middle panel, listed at right side). MOLM-13 cells were infected with shSOCS2 or control lentivirus, selected with puromycin for 48 hours. **(D)** The knockdown of SOCS2 at mRNA level was validated by qRT-PCR. **(E)** ERAD activity evaluated by flow cytometry. Cells were infected with PEZ-NHK, an ERAD reporter lentivirus, and treated with the JAK2 inhibitor Ruxolitinib for 24 hours. **(F)** $ERAD^{high}$ and $ERAD^{low}$ $lineage^{-}$ $c\text{-}Kit^{+}$ $sca\text{-}1^{+}$ LSKs were sorted and then treated with IL-3 (10ng/mL), TPO (10ng/mL), SCF (10ng/mL) for 10min at 37°C.p-STAT5 level was detected by western blot. **(G-I)** MOLM-13 **(G)** and HEL **(H,I)** cells were infected with PEZ-NHK and treated with the JAK2 inhibitor Ruxolitinib **(G, H)** or Fedratinib **(I)** for 24 hours. ERAD activity was evaluated by flow cytometry. **(J)** MOLM-13 cells were infected with PBE2-NHK-Flag lentivirus, selected with Blasticidin S for 48 hours, and treated with Fedratinib (0.4 μM, 24 hours) and CHX (50 uM for 0, 3, 6 hours). Protein level of p-JAK2 and NHK was assessed by western botting. **(K)** HEK 293T cells were transfected with constructs that expressed FLAG-tagged Hrd1, Myc-tagged JAK2-WT, Myc-tagged JAK2-V617F, Myc-tagged JAK2-Y1007/1008F. Hrd1 proteins were immunoprecipitated using FLAG antibody, and JAK2 isoforms binding to Hrd1 were detected by western blot. FACS-purified $SOCS2^{high}$ and $SOCS2^{low}$ HSCs from CD45.2 $SOCS2^{3Xmcherry/+}$ mice were transplanted into irradiated receipt mice along with 0.5 million CD45.1 whole bone marrow competitors. **(L-O)** The contribution of CD45.2 cells in total $CD45^{+}$**(L)**, B ($B220^{+}$; **M**), T ($CD3^{+}$; **N**) and myeloid ($Mac1^{+}$; **O**) cells from peripheral blood was analyzed every 4 weeks for 16 weeks. Data represent mean±s.d. from three independent experiments. Statistical significance was determined using unpaired two-tailed Student's t-test.
